# IGFBPs define distinct pro-tumorigenic CAF subtypes in lung cancer tumour-microenvironment

**DOI:** 10.64898/2025.12.03.692056

**Authors:** Ghanapriya Devi Yengkhom, Kunal Khemraj Nandgaonkar, Sai Kishore Ramanathan, Jan A. Schlegel, Meng Dong, Rahul Thorat, Rajiv Kumar Kaushal, Subhash Yadav, Kumar Prabhash, SharathChandra Arandkar

## Abstract

Cancer-associated fibroblasts (CAFs) play a significant role in influencing tumour outcomes; however, their heterogeneity and context-sensitive nature present a therapeutically challenging target. Here, we identify a group of Insulin-like growth factor binding proteins (IGFBPs), specifically IGFBP5, 6 and 7, as key regulators of CAF heterogeneity in lung cancer. Notably, these binding proteins are highly expressed in lung CAFs and independently modulate CAF phenotype. We demonstrate that tumour-derived cues, particularly TGFβ1 and IL6, affect IGFBP expression in CAFs. Transcriptome studies indicate that selectively knocking down these proteins directs CAFs towards different CAF subtypes. Functionally, CAFs with high levels of IGFBP5, 6, or 7 promote tumour growth in both ex vivo and in vivo settings and, critically, confer cisplatin resistance to tumour cells. Furthermore, the gene signatures associated with these proteins correlate with unfavourable prognosis. In conclusion, our study highlights the phenotypic plasticity of CAFs driven by these three distinct IGFBPs from the same family, consequently influencing the tumour microenvironment landscape.

## Introduction

The importance of the tumour microenvironment (TME) and its components in determining the fate of the tumour cells is now well established in the field. Over the recent years, multiple studies have reported that cancer-associated fibroblasts (CAFs) represent an abundant proportion amongst other stromal cells in most solid tumours (1). Notably, CAFs are known to have an enhanced secretome, migration, invasion and play an important role in tumour progression. On the other hand, certain CAFs have shown a tumour-suppression function, highlighting their heterogeneity and dynamic nature. With advances in single-cell technologies, CAFs have been successfully categorised into different subpopulations, such as myofibroblast, inflammatory, and antigen-presenting CAFs, with each subpopulation having a distinct functional role in mediating tumour outcome and prognosis (2–4). However, emerging evidence indicates that targeting CAF markers has yielded conflicting results (5, 6), highlighting a critical need to understand the mechanisms that drive heterogeneity and their contributions to tumorigenesis. In this study, we have identified Insulin-like Growth Factor Binding Proteins (IGFBPs), a group of secreted proteins, as one of the differentially expressed genes between lung cancer patient-derived normal fibroblasts (NFs) and CAFs. These IGFBPs are known to regulate IGF signalling by sequestering IGF molecules from binding to IGF-I and II receptors and prolonging the half-life of IGF molecules (7, 8); however, some of the IGFBPs mediate IGF-independent functions and exhibit context-dependent roles in multiple tumours (9–11). Recent reports have shown that these proteins also exist in the tumour stroma and, by contrast, have been shown to both sensitise and confer resistance to tumour cells (12–14). This functional duality underscores the need to define their specific roles in the tumour stroma, especially in CAFs.

Here, we elucidate the role of IGFBP5, 6, and 7 in specifying functionally distinct CAF subpopulations. We demonstrate that these IGFBPs regulate CAF cell-autonomous properties, and their expression is modulated by tumour-derived IL6 and TGFβ1. Transcriptomic analysis shows that individually knocking down IGFBP5, 6 or 7 shifts CAFs towards distinct subtypes. Functionally, CAFs with high expression of IGFBP5, 6, or 7 impact tumour cell properties and confer cisplatin resistance. Using immunocompromised mouse models, we confirmed the critical role of these proteins in shaping the tumour outcome. Clinically, the gene signatures derived from IGFBP-positive CAFs are associated with poor prognosis in non-small cell lung cancer (NSCLC) patients. Collectively, our work establishes IGFBP5, 6 and 7 as one of the key mediators of CAF heterogeneity and stromal-driven tumour progression, highlighting a novel stromal axis for therapeutic intervention.

## Results

### IGFBP5, 6, and 7 are differentially expressed in lung cancer-derived CAFs

Previous reports have identified gene signatures that are differentially expressed between normal and cancer-associated fibroblasts (20, 21). To understand the mechanistic insights of CAFs in TME, we re-analysed the RNA-sequencing data of lung cancer patientderived normal fibroblast (NF) and cancer-associated fibroblast (CAF) (16), and identified certain members of Insulin-like Growth Factor Binding Protein (IGFBPs), specifically *IGFBP5, 6,* and *7*, as one of the most differentially expressed genes **(Supplementary Fig. 1a).** To check the expression of IGFBPs in CAFs and other stromal cells, we analysed lung cancer patientderived single-cell RNA-sequencing data using the Curated Cancer Cell Atlas database (22) and observed that *IGFBP5, 6* and *7* are mainly expressed in fibroblasts. Along with fibroblasts, some of these IGFBPs are expressed in pericytes and endothelial cells as well (**Fig. 1a).** We further confirmed the upregulation of IGFBPs at both the cellular mRNA and protein levels in the lung cancer patient-derived NFs and CAFs 4731 **(Fig. 1b & c)**. As IGFBPs are secretory in nature, we collected the conditioned medium from NF and CAF grown in serum-free media to precipitate the secreted proteins. Notably, we observed a clear increase in the secretory levels of IGFBP5, 6 and 7 in CAF compared to NF **(Fig. 1d).** Also, we validated their expression in other patient-derived CAF cell lines (KSH CAF 1054 and CAF 8163) **(Supplementary Fig. 1b).** Though IGFBP5, 6 and 7 are upregulated in CAF 4731 as compared to its counterpart NF, there is a differential expression of IGFBP levels in other patient derived CAF cell lines, suggesting the heterogeneity of these proteins’ expression. Likewise, a similar trend of higher IGFBP expression in fibroblasts is observed in other cancers as well, indicating that their expression is not restricted to lung cancers **(Supplementary Fig. 1c)**. To extend our findings to a larger cohort, we explored two previously published NSCLC single-cell RNA sequencing data (NSCLC_EMTAB6149 and NSCLC_GSE127465) using TISCH (Tumour Immune Single-cell Hub2) (23). Consistently, we observed that all three IGFBPs are majorly expressed in fibroblasts in both datasets. While most fibroblast cells exhibit expression of *ACTA2* (*ACTA2*: αSMA, alpha-smooth muscle actin, classic CAF marker) and IGFBP5, 6, and 7 together, there exist discrete populations of CAF that are *ACTA2*negative but positive for these IGFBPs, again highlighting the heterogeneous dynamics of CAF **(Supplementary Fig. 1d &e)**.

**Fig. 1.**
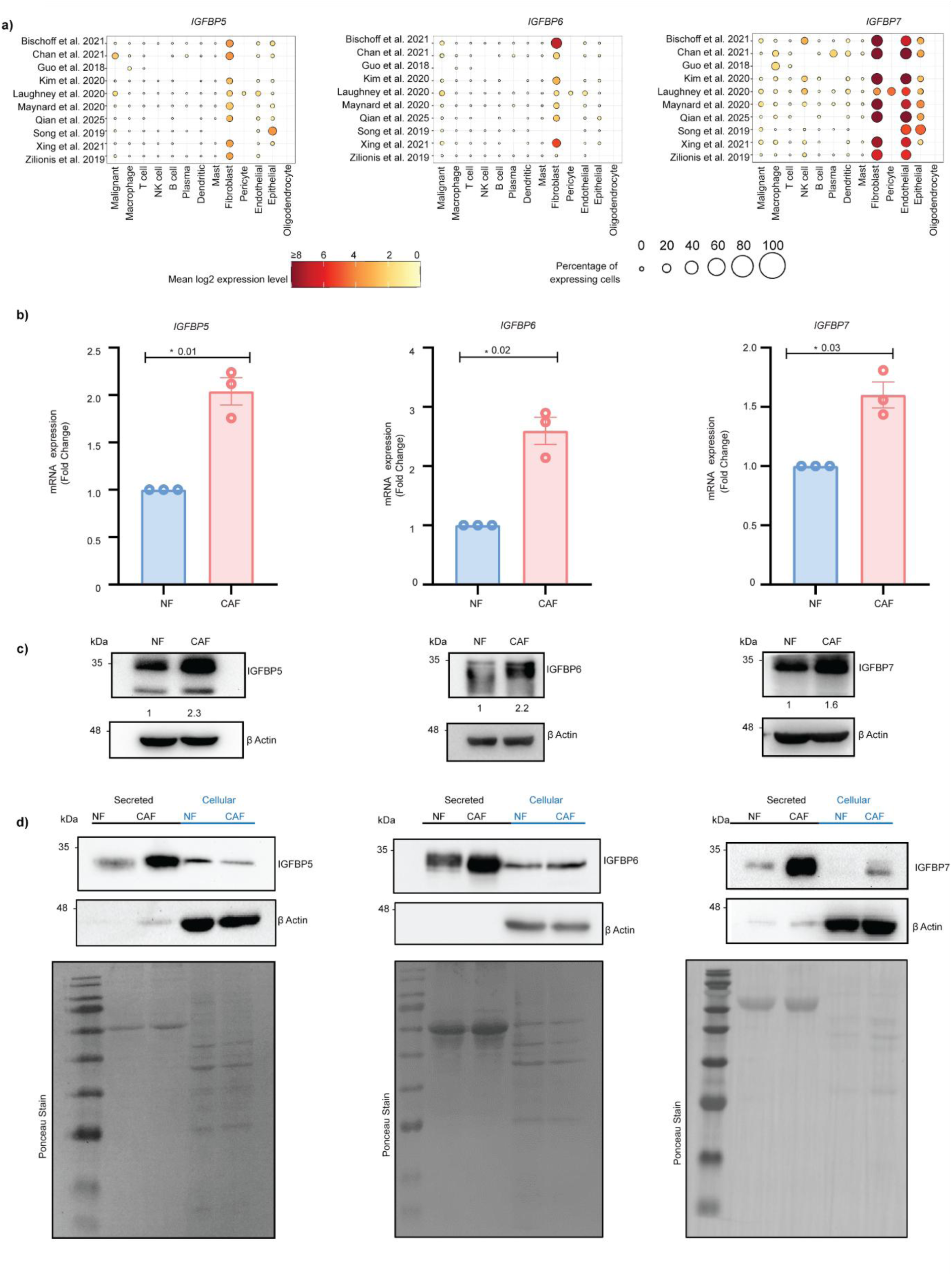
IGFBP5, 6 and 7 are upregulated in CAFs in lung cancer. **a)** Dot plot representing IGFBP5 (left), IGFBP6 (centre) and IGFBP7 (right) expressions in different cell types in publicly available lung single-cell RNA sequencing datasets. Data was generated from the Curated Cancer Cell Atlas. The size of the circle dot represents the percentage of expressing cells, while the colour represents the mean log2 expression level, as indicated in the scale bar (below), **b)** mRNA expression (fold change) of IGFBP5, 6 and 7 in patient-derived lung normal fibroblasts (NF) and cancer-associated fibroblasts (CAF). Data are presented as mean ± SEM (n=3 biological triplicates). Statistical analysis was conducted using an unpaired t-test with Welch’s correction; exact p-values are represented in the graph, **c)** Immunoblots (IBs) showing IGFBP5, 6 and 7 expressions in whole cell lysates of NF and CAF. β-Actin served as the loading control. Densitometric analysis was done using Image Lab (v6.1.0), **d)** IBs showing secreted and cellular (corresponding serum-starved cell lysates) expression of IGFBP5, 6 and 7. β-Actin served as the loading control for serum-starved cellular lysates, while Ponceau blot served as the loading control for secretory lysates. All IBs are representative of three biological triplicates.

### IGFBP5, 6, and 7 are important for maintaining the CAF phenotype

To investigate the cell-autonomous role of IGFBPs, we performed both transient and stable knockdowns of IGFBP 5, 6, and 7 in CAFs **(Fig. 2a)** and validated the knockdown efficiency at both mRNA **(Supplementary Fig. 2a)** and protein level **(Fig. 2b & Supplementary Fig. 2b).** While it was previously reported that these proteins exhibit functional redundancy (24), we observed no significant compensation by the other IGFBP proteins under individual knockdown **(Supplementary Fig. 2a).** Interestingly, we observed that each IGFBP knockdown exhibited a different cellular morphology in terms of actin arrangement pattern, suggesting that these proteins may influence the cytoskeleton dynamics of CAFs **(Fig. 2c & d).** In TME, CAFs display an activated fibroblast phenotype, which can be assessed by the collagen contraction assay. Furthermore, we observed that individual knockdown of IGFBP5, 6, and 7 leads to reduced contractility, while triple knockdown (Pooled knockdown) leads to a complete loss of contractile property of fibroblasts **(Fig. 2e-g).** Moreover, a distinct difference was observed in the levels of key actomyosin regulators, phosphorylated myosin light chain (p-MLC) **(Supplementary Fig. 2c-e)**. A similar trend of loss of contractility was observed in KSH1054 CAF cells **(Supplementary Fig. 2f-h),** indicating that these binding proteins maintain the contractility and assembly of stress fibres in CAFs.

**Fig 2.**
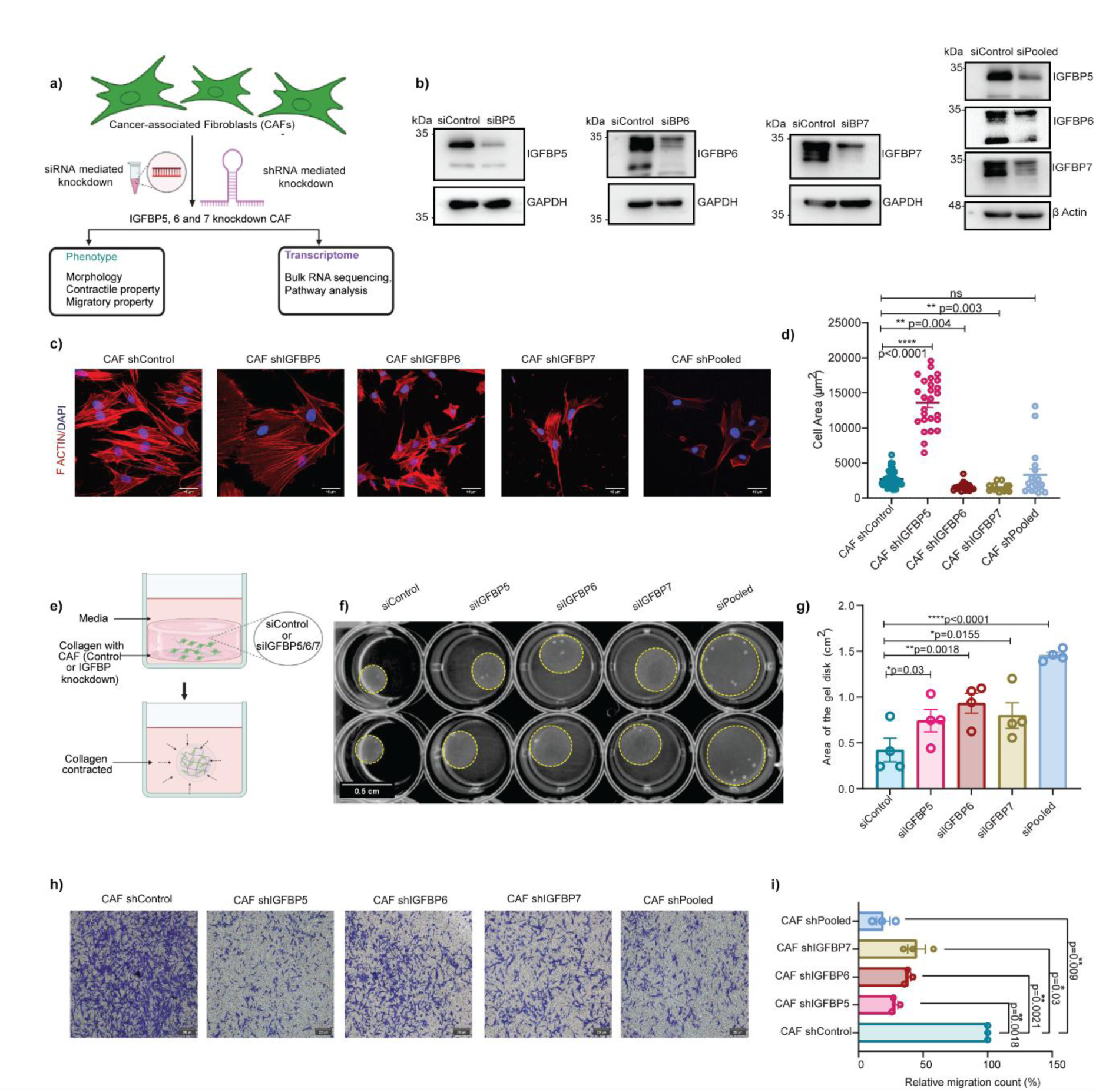
IGFBP5, 6 and 7 regulate CAF’s properties. a) Schematics showing investigation of IGFBP5, 6 and 7 roles in CAF biology through transient and stable knockdown approaches. Illustration was prepared using Biorender.com, **b)** Immunoblots showing knockdown validation of IGFBP5, 6 and 7 in CAF upon transient transfection. GAPDH and β-Actin served as the loading control. The IBs represent three biological replicates, **c)** Representative immunofluorescence images showing F-actin staining (red) and DAPI (blue) in CAF upon individual and pooled knockdown of IGFBPs, scale bar 45μm, **d)** Quantification of cell area for **(c).** Quantification was performed for a minimum of thirty cells, which were included from three biological replicates. Data are presented as mean ± SEM, with open circles indicating individual cells. Statistical analysis was conducted using the Kruskal-Wallis test with Dunn’s multiple comparison; exact p-values are indicated in the graph, **e)** Schematics showing collagen contraction experiment. Illustration was prepared using Biorender.com, **f)** Image representing technical duplicates of collagen contraction upon individual and pooled knockdown of IGFBP5, 6 and 7. Yellow dotted lines mark the boundary of the gel, **g)** Quantification of the area of the gel disk is represented as a bar plot with mean ± SEM, n=4 biological replicates, **h)** Representative brightfield images of migrated cells (crystal violet stained) after IGFBP5, 6 and 7 knockdowns, scale bar 200μm. Images were captured 16 hours after wiping off the non-migrated cells in the upper chamber, **i)** Cell count of the migrated cells is taken as an average of four different fields of one experiment. Data represent mean (relative cell count with respect to control (%) ± SEM, n= 3 biological replicates, **(g, i)** To determine statistical significance, One-way ANOVA with Dunnett’s multiple comparison was used; exact p-values are indicated in the graph.

CAFs are known to have an enhanced migratory property in the TME towards tumour cells or tumour cell-secreted factors (25). To investigate the IGFBP-mediated CAF migratory properties, we performed a trans-well migration assay where CAFs with different IGFBP knockdowns were seeded in the upper chamber, and serum in the medium was used as a chemoattractant. We observed that the knockdown of individual IGFBPs reduced the migratory property of CAF, with triple knockdown of these proteins exhibiting a pronounced decrease in migration, indicating an additive effect of IGFBP5, IGFBP6, and IGFBP7 in maintaining CAF phenotype (**Fig. 2h & i).**

### IGFBPs regulate different CAF-subtype gene signatures

Multiple single-cell RNA sequencing studies revealed that different CAF subtypes exist across different cancers, with each subtype having a distinct function (2, 26, 27). To understand how each IGFBP mediates the rewiring of CAF gene expression, we performed bulk RNA sequencing for siIGFBP5, siIGFBP6, siIGFBP7, siPooled, and siControl (non-targeted) CAF cells. Interestingly, we observed a distinct class of mRNA changes upon individual knockdown.

Upon IGFBP5 knockdown, 1250 genes were upregulated and 1416 genes were downregulated **(Fig. 3a).** To elucidate the enriched pathways, gene set enrichment analysis of the differentially expressed genes upon IGFBP5 knockdown was performed, revealing that pathways such as collagen fibril organisation, interleukin-6 production, and type-II interferon production were upregulated, while most pathways associated with stress fibre assembly were downregulated **(Fig. 3d)**. Importantly, *MACIR*, known to regulate M2 macrophage functions (28), was down-regulated upon IGFBP5 knockdown **(Supplementary Fig. 3a)**, while *CXCL10* was up-regulated, implying that CAF-derived IGFBP5 may be involved in modulating the synergistic effect with macrophages in influencing tumour outcome **(Supplementary Fig. 4d).** To further decode which CAF subtype IGFBP5 regulates, we compared the gene expression with the known CAF-subtype gene markers from Curated Cancer Cell Atlas **(Supplementary Table 2).** Surprisingly, we observed that IGFBP5 knockdown leads to upregulation of MHCII, iCAF and myCAF gene signatures **(Fig. 3g).** In line with the available single-cell RNA sequencing-generated databases where IGFBP5+ fibroblast is associated with PI16+ gene signatures **(Supplementary Fig. 4g)**, our findings also suggest that IGFBP5 knockdown leads to the downregulation of PI16+ CAF gene signatures. Additionally, we validated some of the known CAF markers and observed that *ACTA2, IL6* and *CXCL10* are upregulated at the mRNA level, inferring that IGFBP5 regulates both myCAF and inflammatory-associated genes **(Supplementary Fig. 4d).** Upon IGFBP6 knockdown, 2362 genes were upregulated and 2576 genes were downregulated **(Fig. 3b).** Gene set enrichment analysis revealed that lipid transport, inflammatory response, and xenobiotic metabolism were upregulated and DNA replication pathways were downregulated **(Fig. 3e).** Certain genes related to E2F targets and G2M checkpoints, such as *CDC45, E2F8 and DARS2*, were downregulated upon IGFBP6 knockdown **(Supplementary Fig. 3b)** suggesting their role in shaping a dividing-like CAF phenotype. Emerging evidence highlighted that certain CAF gene clusters enriched in metabolism-related pathways exist in different cancers (29). Interestingly, IGFBP6 is also involved in pathways related to xenobiotic metabolism **(Supplementary Fig. 3c).** From further analysis, we found that IGFBP6 regulates lipid metabolism, complement, pericyte-like, and stress CAF subtype-associated gene signatures **(Fig. 3h).** In line with this, we observed that IGFBP6-expressing CAFs in the lung show low expression of pericyte-like and lipid metabolism gene signatures **(Supplementary Fig. 4h).** Additionally, we observed that myCAF-associated gene signatures are upregulated upon IGFBP6 knockdown **(Supplementary Fig. 4e).** Overall, this suggests that IGFBP6 is involved in regulating metabolism as well as inflammatory responses in CAF.

**Fig 3.**
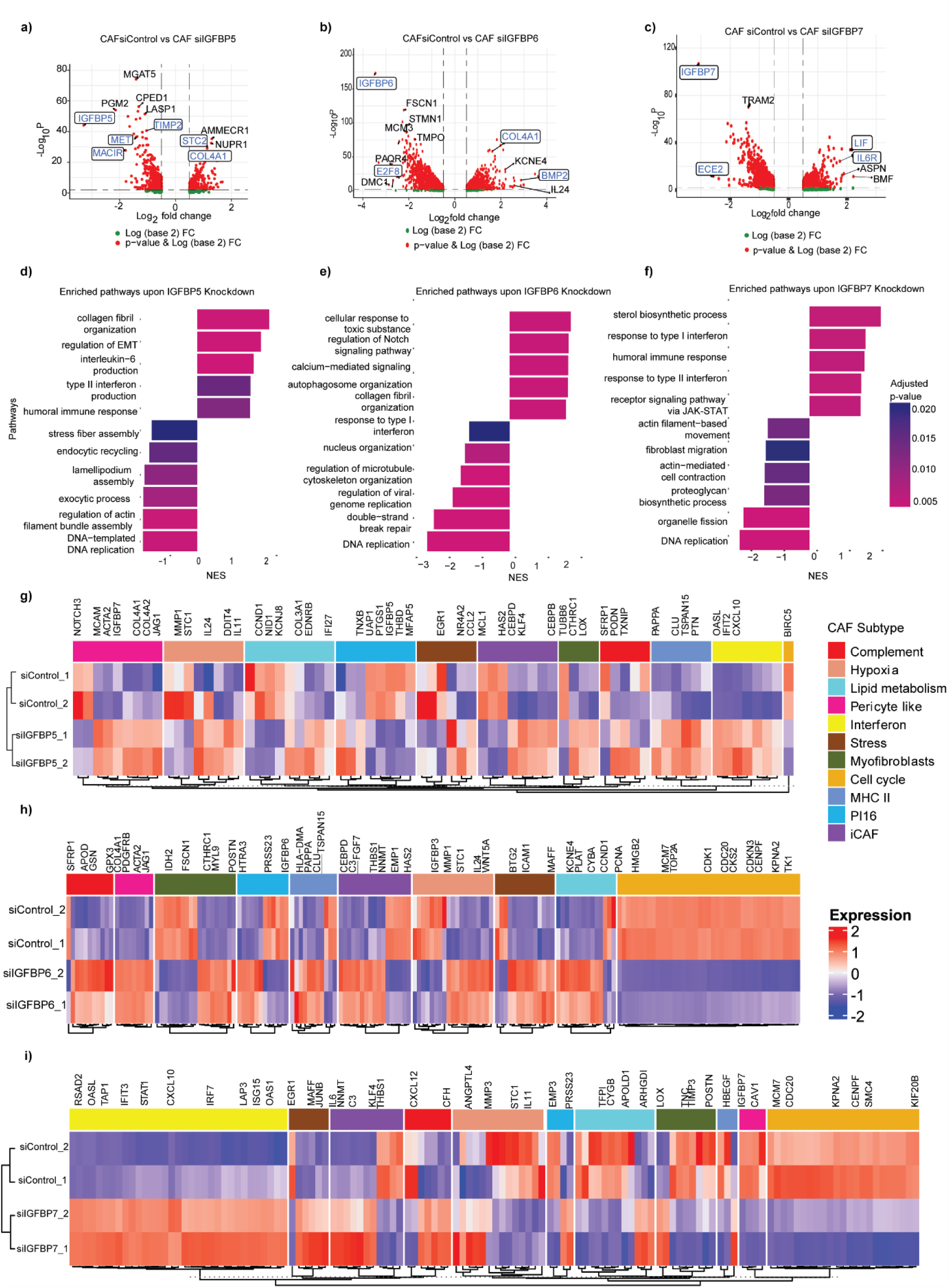
IGFBP knockdown directs CAF to distinct subtypes. **a)** Volcano plot representing differentially expressed genes between CAF siControl vs siIGFBP5 **b)** CAF siControl vs siIGFBP6 **c)** CAF siControl vs siIGFBP7 (n=3 biological replicates). A log2 fold change of 0.5 was set as the threshold. Selected genes for validation are marked in blue. **d)** Enriched pathways based on the differentially expressed genes upon IGFBP5 knockdown **e)** IGFBP6 knockdown **f)** IGFBP7 knockdown **g)** Heatmap comparing the published CAF subtype gene sets with CAF siControl versus siIGFBP5 **h)** CAF siControl versus siIGFBP6 **i)** CAF siControl versus siIGFBP7 (n=2 biological replicates)

On the other hand, IGFBP7 knockdown revealed 2045 upregulated and 2368 downregulated genes **(Fig. 3c).** The pathways associated with interferon responses and immune activation were upregulated under IGFBP7 knockdown, while migration and contraction-related pathways were downregulated **(Fig. 3f).** Genes such as *TAPBP*, involved in antigen presentation and T-cell activation (30), were enriched, suggesting that IGFBP7-expressing CAF may also be involved in modulating antigen presentation and immune infiltration. Also, genes related to microtubule cytoskeleton organisation and collagen type I synthesis pathways (31), such as *TRAM2* were found to be down-regulated at mRNA levels **(Supplementary Fig. 3d).** Additionally, *LIF,* which is a known iCAF marker, was found to be highly upregulated upon IGFBP7 knockdown, suggesting that IGFBP7 may be involved in regulating the transition of CAFs to inflammatory CAFs **(Supplementary Fig. 4f).** Consistently, we noticed a similar trend where interferon CAF subtype-associated gene signature was upregulated along with iCAF in IGFBP7 knockdown CAF; on the other hand, myCAF, pericyte-like and cell-cycle CAF subtype-associated gene signatures were downregulated **(Fig. 3i, Supplementary Fig. 4i).** We validated the increased expression of interferon CAF markers at mRNA level **(Supplementary Fig. 4f).** Notably, upon triple knockdown of IGFBP5, 6 and 7, DNA replication pathway was downregulated while metabolic, inflammatory and collagen organization pathways were upregulated **(Supplementary Fig. 4a & b).** Additionally, on comparison with the known CAF subtype gene signatures, iCAF, lipidmetabolism, pericyte-like, and MHC-II CAF subtypes were upregulated **(Supplementary Fig. 4c).** Collectively, 162 genes were commonly regulated by all the IGFBP5, 6, 7 and triple knockdown, which mainly regulate cell proliferation, migration and stress response **(Supplementary Fig. 4j).**

Although there exists a small degree of similarity in terms of functional phenotypes and pathways, interestingly, each IGFBP has its own independent mechanism of modulating CAF phenotype, and all IGFBP5, 6 and 7 have a role to play in defining CAF heterogeneity.

### TGFβ and IL-6 regulate IGFBPs expression in CAF

Different cues derived from tumour cells direct CAFs to behave differently and govern CAF heterogeneity (32). To dissect the mechanism of how IGFBPs regulate the CAF phenotype, we utilised NicheNet (19), a computational method that infers the link between active ligand and target genes. To this end, we selected a set of defined ligands based on the existing literature (33), which have the potential to influence the tumour microenvironment. To define the ligands, we selected only those ligands which are expressed by lung tumour cells and whose cognate receptors are expressed on the target CAF cells. We observed that VEGFA, FGF2, EGF and TGFβ1 had the highest ligand activity potential in regulating the IGFBP5, 6 and 7 differentially expressed genes, respectively **(Fig. 4a, d & g).** To further investigate, we performed active ligand target analysis using the differentially expressed genes as our target. Out of all the potential ligands regulating IGFBP5 differentially expressed genes, we found that TGFβ1, FGF2 and IL6 can regulate IGFBP5 expression **(Fig. 4b).** Next, we used NicheNet’s weighted integrated signalling and gene regulatory networks to directly visualise the signalling interaction between our ligands of interest and IGFBP5. We observed that while TGFβ1 can directly interact with IGFBP5, IL6 and FGF2 can regulate IGFBP5 expression through STAT3 or MAPK3 **(Supplementary Fig. 5a).** To test this, we directly treated the CAF with recombinant IL6 and TGFβ1 for 48 hours **(Supplementary Fig. 5d-f).** We found that IGFBP5 expression was increased with IL6 treatment, while treatment with TGFβ1 led to IGFBP5 protein downregulation **(Fig. 4c).** Our observations are consistent with the previous findings, where TGFβ1 inhibits IGFBP5 (34) and IL6 can induce IGFBP5 expression in senescence-induced fibroblasts (35). It is reported that TGFβ signalling regulates myCAF phenotype while IL-1α induces an inflammatory phenotype leading to IL6 expression(36). Interestingly, we observe that upon IGFBP5 knockdown, both αSMA and IL-6 are upregulated, implying that IGFBP5 plays a role in regulating myCAF and iCAF phenotype up to some extent **(Fig. 4c).** On a similar note, we observed that IL6 treatment upregulates IGFBP6 expression while it is downregulated upon treatment with TGFβ1 **(Supplementary Fig. 5d-f).** Additionally, upon IGFBP6 knockdown, pSTAT3 is further reduced along with IL-6, suggesting that the STAT3-IL6 axis is involved in regulating IGFBP6 expression in CAF **(Fig. 4f).** Similar to the IGFBP5 knockdown, we also observed an increase in αSMA levels, but a reduction in IL-6 levels, implying that IGFBP6 positively regulates iCAF phenotype while restraining myCAF phenotype to an extent **(Fig. 4c)**. Importantly, IL6R is also upregulated when IGFBP7 is knocked down **(Supplementary Fig. 3d.),** leading to an increase in IL6 expression, implying that IGFBP7 restrains inflammatory CAF **(Fig. 4i).** Though IGFBP5, 6 and 7 are upregulated in CAF, it is intriguing to observe how these proteins respond to different tumour microenvironmental cues and mediate CAF dynamics accordingly.

**Fig. 4.**
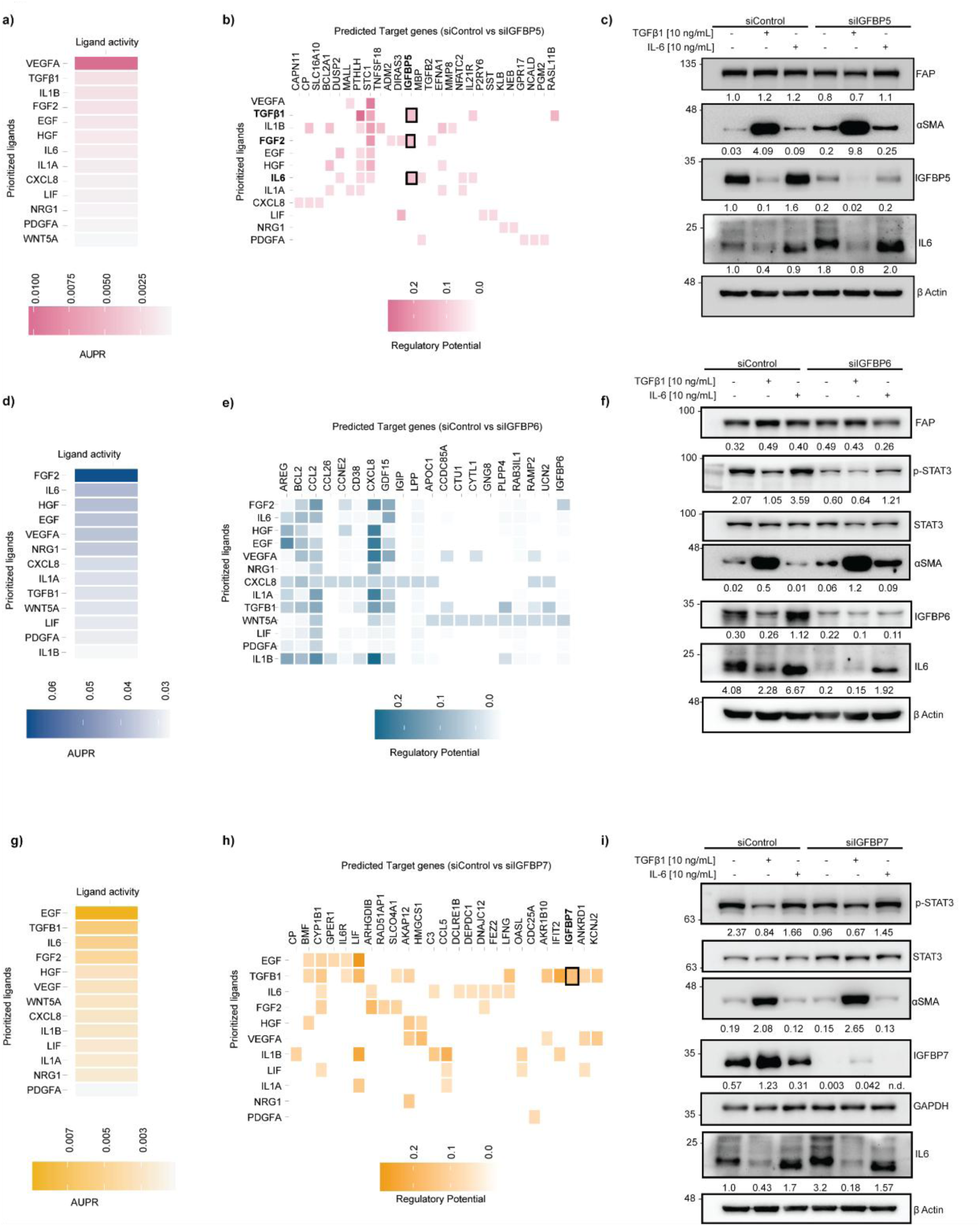
TGFβ1 and IL6 regulate IGFBP expression in CAF. **a)** Ligand activity analysis showing the list of potential ligands that can regulate differentially expressed genes upon IGFBP5 knockdown in CAF, **d)** IGFBP6 knockdown, **g)** IGFBP7 knockdown, **b)** Ligand-target matrix showing the regulatory potential of potential ligands with CAF siControl vs siIGFBP5, **e)** CAF siControl vs siIGFBP6, **h)** CAF siControl vs siIGFBP7, **c)** Western blot analysis of CAF siControl and siIGFBP5, **f)** CAF siControl and siIGFBP6, **i)** CAF siControl and siIGFBP7, after treatment with TGFβ1[10 ng/mL] and IL6 [10 ng/mL] for 48 hours. Densitometric analysis was done using Image Lab (v6.1.0). All IBs are representative of three biological triplicates.

We next investigated the IGFBP-mediated IGF signalling axis in CAFs. We observed that upon transient knockdown of IGFBP5, 6 and 7, there is a reduction in p-ERK and p-Akt levels despite a slight increase in the levels of pIGF1R, suggesting that these IGFBPs may have an IGF-independent function in rewiring CAF functions **(Supplementary Fig. 6a-c).** To further delineate what other pathways IGFBPs regulate, we used decoupleR to infer pathway activities **(Supplementary Fig. 6d-f)** (37). In common, we observed that the PI3K and MAPK pathway is downregulated upon IGFBP5, 6 and 7 knockdowns. Upon IGFBP5 knockdown, the TGFβ pathway is upregulated with NFκB and JAK-STAT. On the other hand, TGFβ, NFκB, and JAK-STAT pathways are downregulated with IGFBP6 knockdown. Similar to IGFBP5, we observe that p53, JAK-STAT and NFκB are upregulated upon IGFBP7 knockdown. Altogether, other than the IGF-dependent axis, clearly, IGFBP5, 6 and 7 can differentially respond to TME cues and regulate the CAF biology by utilising multiple and diverse signalling pathways.

### High IGFBP-expressing CAFs can influence tumour growth

To understand the influence of IGFBP-expressing CAFs on tumor cell behaviour, we treated H460 tumor cells with conditioned medium (CM) derived from CAF-stable IGFBP5, IGFBP6, IGFBP7 and Pooled knockdown cells. As expected, we observed a clear increase in tumour cell proliferation with CM derived from CAF shControl cells. Strikingly, our results suggest that CAF IGFBP5, 6 and 7 control tumor cell proliferation non-autonomously **(Fig. 5a & b)**. Interestingly, we also observed a decrease in tumour cell migration **(Supplementary Fig. 7a & b)** and invasion properties upon IGFBPs knockdown in CAFs **(Fig. 5c-f).** It is already known that CAFs contribute to chemoresistance to various drug molecules in multiple cancers (38–40). To investigate the role of CAF-derived IGFBPs in chemoresistance, we treated H460 tumour cells with cisplatin along with the conditioned media derived from different IGFBP knockdown CAF cells. We observed that IGFBP-expressing CAFs can confer cisplatin resistance **(Fig. 5 g & h**). Together, this suggests that each IGFBP has a unique signature and can exert non-autonomous function on tumour cell properties.

**Fig. 5.**
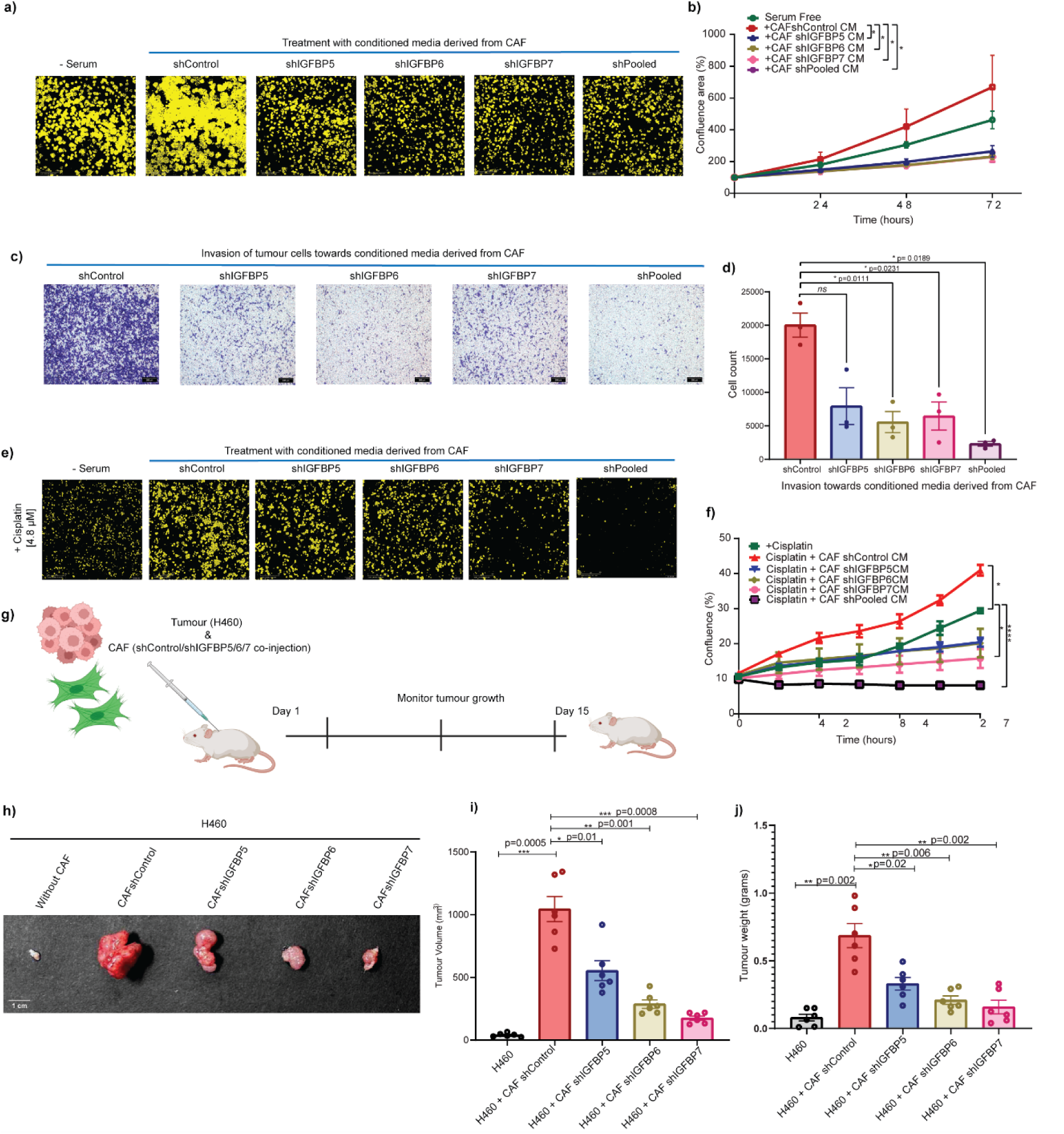
IGFBP-expressing CAF impacts tumorigenesis ex vivo and in vivo. **a)** H460 cells (masked-yellow) treated with conditioned media (CM) derived from CAF individual and pooled knockdowns of IGFBP5, 6 and 7, image shown here is a representative for the 72-hour timepoint, **b)** Graph represents mean ± SEM showing the proliferation of tumour cells represented as confluence percentage over a 72-hour time interval, the line colour represents different conditions, n=3, * indicates p-value less than 0.05, **c)** Representative brightfield images of invaded cells (crystal violet stained) of tumour cells towards conditioned media derived from individual and pooled IGFBP5, 6 and 7 knockdowns in CAF, scale bar 200μm. Images were captured 16 hours after wiping off the non-invaded cells in the upper chamber. **d)** Cell count of the invaded cells is taken as an average of four different fields of one experiment. Data represent mean ± SEM, n=3 biological replicates. To determine statistical significance, Oneway ANOVA with Dunnett’s multiple comparison was used; exact p-values are indicated in the graph, **e)** H460 cells (masked- yellow) treated with the IC_50_ values of cisplatin [4.8 μM] with CM derived from CAF individual and pooled knockdowns of IGFBP5, 6 and 7, image shown here is a representative for the 72-hour time-point, **f)** Graph represents mean ± SEM showing the tumour cells represented as confluence percentage over a 72-hour time interval, the line colour represents different conditions, n=3, * indicates p-value less than 0.05, **** indicates pvalue less than 0.0001. **g)** Schematics showing in-vivo experiment design, illustration was prepared using Biorender.com. **h)** m-Cherry expressing H460 tumour cells were injected subcutaneously alone (0.5 * 10^6^ cells) or co-injected with GFP-expressing CAF individual IGFBP5, 6 or 7 knockdown or shControl (1.5 * 10^6^ cells) in male NOD-SCID mice. Tumours were excised on the 16^th^ day. **i)** Tumour volume and **j)** tumour weight were measured after the tumours were excised; data are represented for 6 mice from three independent experiments. Dunnet’s multiple comparison test was used to determine statistical significance.

Multiple studies have shown that CAFs promote tumorigenesis (16, 41). To investigate the contribution of these IGFBP-expressing CAFs on tumorigenesis in vivo, we co-injected GFP-labelled CAF shControl or individual shIGFBP5, 6, or 7 with mCherry luciferase-expressing H460 tumour cells in immunocompromised NOD-SCID mice **(Fig. 5g).** The tumour volume was monitored at regular intervals for fifteen days. In agreement with the previous literature, CAFs support tumour growth compared to tumour cells alone. Interestingly, individual IGFBP knockdown in CAFs reduced tumour growth to various degrees **(Fig. 5h-j).** Collectively, these results indicated that CAF IGFBP5, 6 and 7 are important for determining overall tumour outcome.

### IGFBP5, 6 and 7 expressions in the stroma of NSCLC patients are associated with poor patient outcome

To investigate the spatial expression of IGFBPs in NSCLC patient tissues, immunohistochemical and immunofluorescence staining were performed. Consistent with the previous literature (42), we also observed high stroma content in lung adenocarcinoma and squamous cell carcinoma patients, along with high collagen deposition **(Fig. 6a).** We confirmed the expression of IGFBP5, 6 and 7 in stromal fibroblasts in addition to tumour cells. Multiplex immunofluorescence staining revealed the heterogeneity in terms of spatial distribution of IGFBP5, 6, and 7 in αSMA and FAP positive cancer-associated fibroblasts **(Fig. 6b).** Additionally, immunofluorescence of these patient-derived tissues revealed co-staining of αSMA with IGFBP5, 6 or 7; however, not all αSMA-positive CAFs express these IGFBPs, implying their heterogeneous expression **(Supplementary Fig. 7c).** Furthermore, the abundance of distinct CAF subtypes is associated with different survival outcomes in multiple cancers (43). To determine whether IGFBP-expressing CAFs have any correlation with survival outcomes in lung cancer, we curated IGFBP5, 6, and 7-specific gene sets from RNAseq data by including those genes which are positively regulated by each specific IGFBP to estimate survival outcomes using lung adenocarcinoma datasets from TCGA. We observed that patients with high expression of CAF_IGFBP5, 6 and 7 gene signatures are associated with poor survival, while patients exhibiting lower expression of these gene sets are associated with favourable clinical outcomes **(Fig. 6c, d & e).** Collectively, our findings indicate that IGFBP5, 6 and 7 shape the CAF landscape and contribute to determining clinical outcomes in lung cancer **(Fig. 7).**

**Fig. 6.**
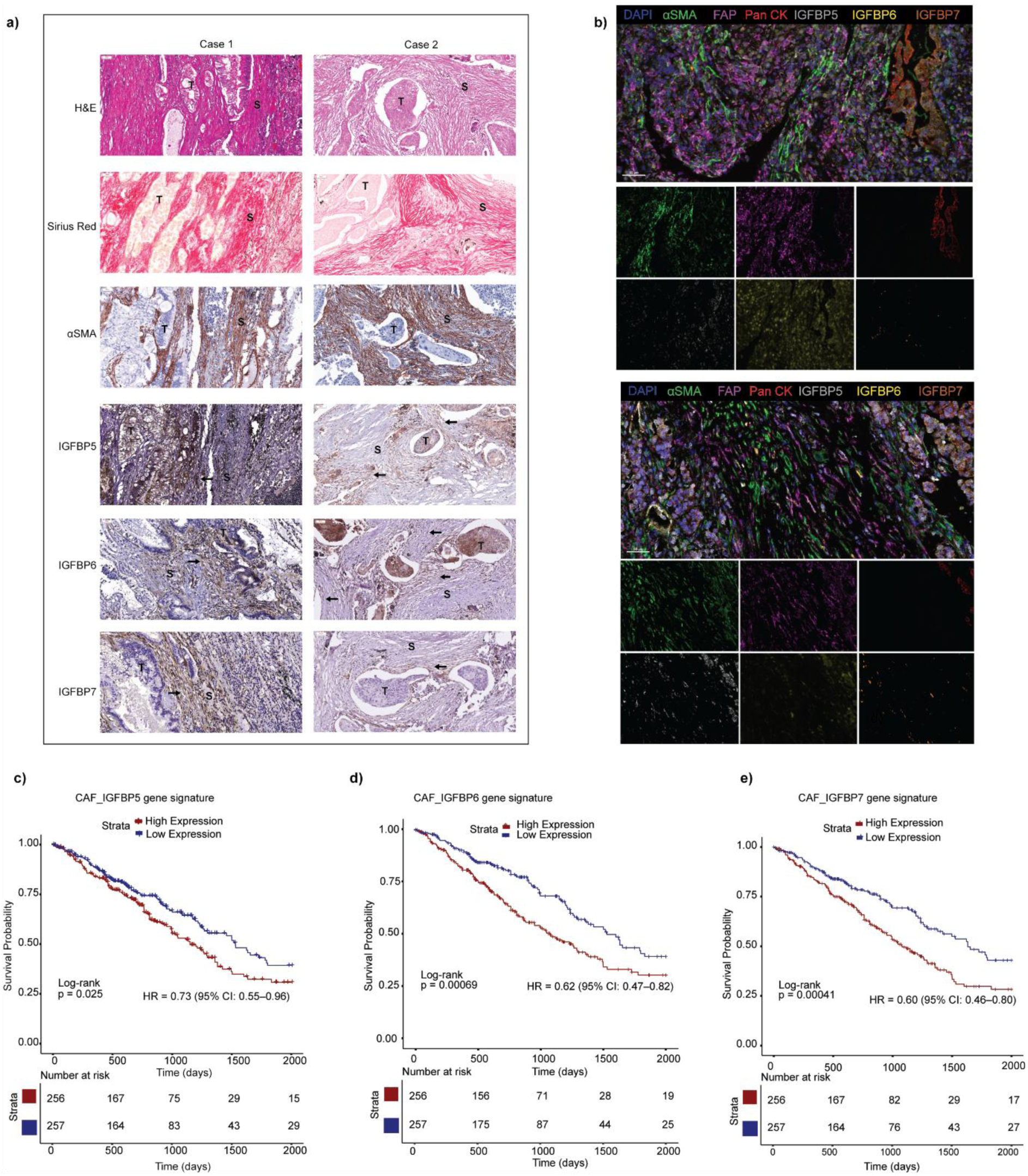
IGFBP5, 6 and 7 are expressed in human lung cancer stroma and correlate with poor survival. **a)** Formalin-fixed paraffin-embedded NSCLC tumour sections were stained with hematoxylin and eosin, Sirius Red, antibodies against α-SMA, IGFBP5, 6 and 7. Cases 1 and 2 represent two different patients. ‘T’ indicates tumour and ‘S’ indicates stroma. The arrow indicates the expression of the respective proteins in fibroblasts; scale bar, 50 μm. **b)** Representative multiplex immunofluorescence staining images of IGFBP5+, IGFBP6+ and IGFBP7+ CAFs. DAPI (blue), αSMA (green), FAP (pink), PanCK (red), IGFBP5 (grey), IGFBP6 (yellow), IGFBP7 (orange) in human non-small-cell lung cancer tissue sections. Scale bars 50 μm. **c)** survival between high and low expression of CAF_IGFBP5 gene signatures **d)** CAF_IGFBP6 gene signatures **e)** CAF_IGFBP7 gene signatures in TCGA-LUAD datasets, pvalues were obtained from two-sided log-Rank tests.

**Fig. 7.**
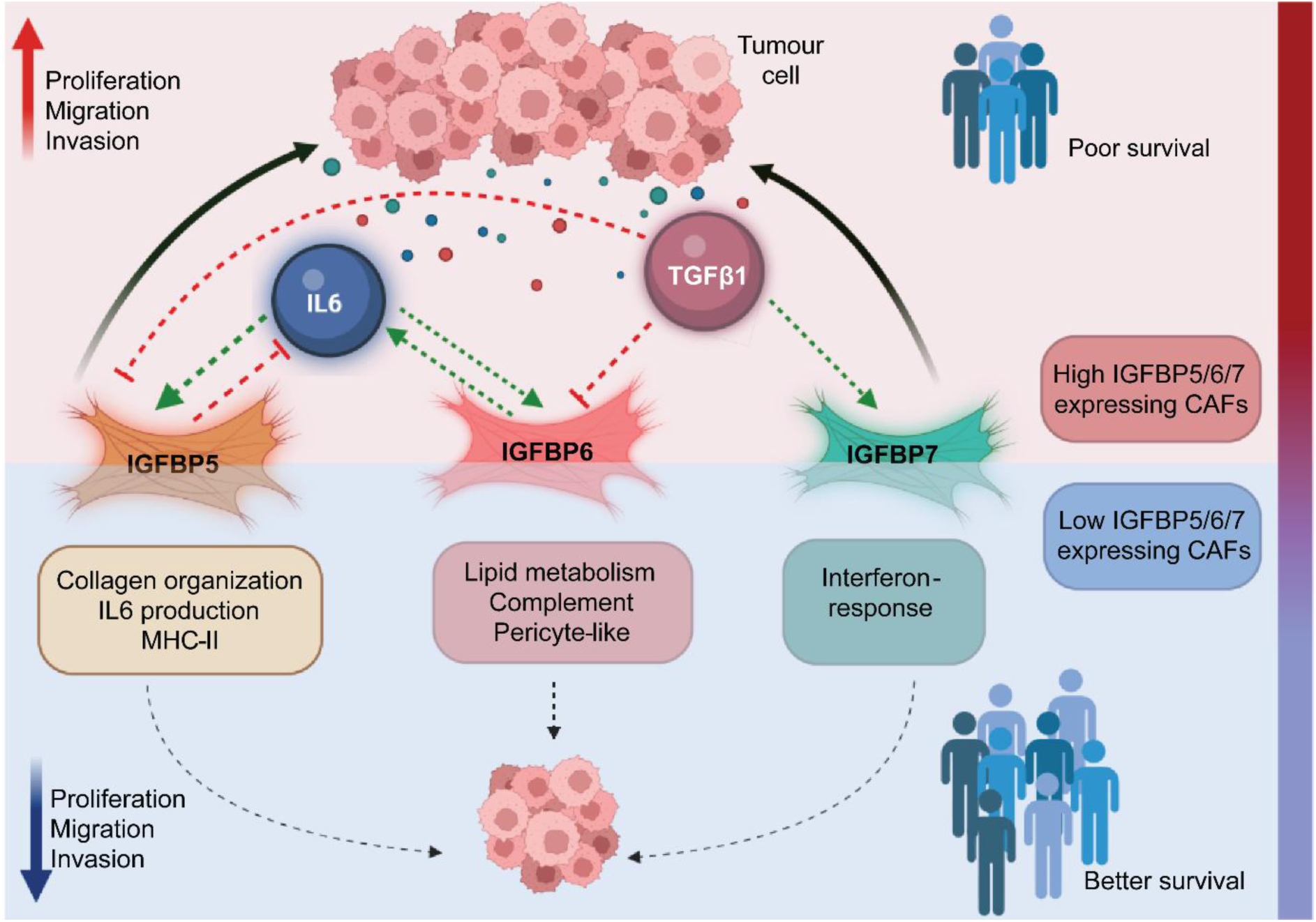
A schematic abstract illustrating the cross-talk of tumour and IGFBP-expressing CAFs. Tumour cells secrete TGFβ1 and IL6, which regulate IGFBP5, 6 and 7 expressions in cancer-associated fibroblasts. These high IGFBP-expressing CAFs show pro-tumorigenic effects and are associated with poor patient outcomes. On the other hand, knocking down IGFBP5, 6, and 7 selectively pushes CAF towards distinct subtypes, which suppresses tumour properties and correlates with better survival.

## Discussion

The functional and phenotypic heterogeneity of cancer-associated fibroblasts presents a major challenge in their therapeutic targeting. Here, we dissect this heterogeneity to convert pro-tumorigenic CAFs into a tumour-restraining state. We identified a group of proteins from the IGFBP family, mainly IGFBP5, 6, and 7, as differentially expressed genes in lung cancer patient-derived CAFs compared to NFs. A previous study by Rix et al. (12) also confirmed that the levels of these three proteins were high in NSCLC-derived CAFs. Notably, these proteins exhibit context-dependent behaviour in different cancers (7, 46); however, their contributions to CAF properties have not been thoroughly investigated. In the present study, we report for the first time that IGFBP5, 6, and 7 regulate CAF phenotype in a cell-autonomous manner by rewiring the transcriptomic landscape, thereby driving them into distinct subtypes. Specifically, we found that IGFBP5 negatively regulates mainly myofibroblasts (myCAF), MHC-II, and inflammatory (iCAF) CAF subtypes. Conversely, IGFBP6 in CAFs reveals a dual role- it positively regulates cell-cycle progression and DNA replication while suppressing lipid metabolism and pericyte-like phenotypes. This aligns with the established roles for IGFBP6, previously associated with lipid metabolism in endothelial cells and DNA replication pathways (47–49). Recent studies have identified interferon-response CAF, a novel CAF subtype in solid cancers with context-dependent roles (26, 50–52). Interestingly, our study links IGFBP7 to interferon signalling in CAFs, revealing a shift towards an interferon-high, inflammatory-high subtype. It has been demonstrated that IGFBP7 also plays a role in macrophage polarisation and poor prognosis in gastric cancers (14). In line with this, we also observed certain immune signature-related genes to be upregulated upon IGFBP7 knockdown, suggesting it mediates CAF-immune cell crosstalk. Investigating this interplay could yield insights for predicting and designing effective immunotherapies.

Within the TME, cancer cells secrete certain signalling molecules to define the surrounding stromal cell types and their heterogeneity (53). We delineated a regulatory network in the TME where tumour-derived IL6 and TGFβ1 drive distinct IGFBP-mediated feedback loops in CAFs. IGFBP6 establishes a positive feedback circuit via the IL6-STAT3IGFBP6 axis, while IGFBP5 feeds back to modulate IL6 production. On the other hand, TGFβ1 upregulates IGFBP7, which in turn, acts as a negative regulator of IL6 and LIF, the key mediators of inflammatory CAFs. Meanwhile, both IGFBP5 and 6 act as negative regulators of αSMA, a well-defined myCAF marker. Clinically, gene signatures for CAF-derived IGFBP5, 6 and 7 are all associated with poor survival in NSCLC patients.

We also addressed the cell-non-autonomous functions of IGFBP-expressing CAFs. Remarkably, our data point towards CAF IGFBP-mediated pro-tumorigenic role in both *in vivo* and *ex vivo* settings. Moreover, it has been demonstrated that tumours with fibrotic stroma impede the availability of drugs, thereby contributing to chemoresistance (38, 39, 54). Intriguingly, we observed that all IGFBP5-, 6-, and 7-expressing CAFs confer cisplatin resistance to tumour cells. Whether this effect pertains to other common chemotherapeutic agents or IGFBP-expressing CAF has a common mechanism against chemotherapies is still unknown.

Collectively, our study emphasised that IGFBP5, 6 and 7 are pivotal for maintaining the CAF phenotype by independently rewiring the transcriptome. Defining the signalling mechanisms and immune interactions of these IGFBP-defined CAF subpopulations will be crucial for predicting patients’ responses to immunotherapy.

## Materials and Methods

### Ethics statement

This study complied with all the relevant ethical regulations. All animal procedures were approved and conducted in strict accordance with the rules and regulations of the Institutional Animal Ethics Committee (IAEC) (Project No. 09/2023). All patient tissue blocks were obtained in accordance with the guidelines of the Institutional Ethics Committee of the Tata Memorial Centre, ACTREC (Project No. 901009).

### Cell culture and reagents

NSCLC tumour cell line H460 was maintained in RPMI with 10% FBS (Fetal Bovine Serum, Gibco South America). Lung cancer patient-derived normal adjacent fibroblasts (NFs) and cancer-associated fibroblasts (CAFs) (4731, KSH CAF1054, CAF 8163) (kind gift from Prof. Moshe Oren’s Lab) were maintained in RPMI (Gibco South America) containing heatinactivated 10% FBS at 37⁰C in an atmosphere of 5% CO_2_. All cell lines were tested for *Mycoplasma* routinely using a Mycoplasma Detection kit (EZ-PCR Mycoplasma, Biological Industries)

### RNA Isolation and RT-qPCR

RNA was isolated using the standard Trizol extraction method (Sigma-Aldrich). cDNAs were synthesised using a cDNA kit (Thermo Scientific). The absolute mRNA expression of genes was measured by qRT-PCR with SYBR-Green Master Mix (TB Green, Takara) and primers **(Supplementary Table 1).**

### Western Blot for whole cell lysates

Cells were lysed directly with RIPA buffer or Laemmli buffer and subjected to SDS PAGE and primary antibodies of α-Smooth Muscle Actin (#19245, CST, 1:1000), FAP (#66562, CST, 1:1000), S100A4 (#13018, CST, 1:1000), IGFBP5 (sc-515184, Santa Cruz, 1:1000), IGFBP6 (AF876, R&D Systems, 1:1000), IGFBP7(sc-365293, Santa Cruz 1:750), Phospho-p44/42 MAPK (Thr202/Tyr204) (#9101, CST, 1:2000), p44/42 MAPK (Thr202/Tyr204) (137F5)(#4695, CST, 1:2000), Phospho-Akt (Ser473) (#4060, CST, 1:2000), Akt (pan) (#2920, CST, 1:2000), Phospho-Insulin R(Y1162/Y1163)/IGF-1R (Y1132/Y1136) (AF2507, R&D Systems, 1:1000), IGF-I R/IGF1R (MAB391, R&D Systems, 1:1000), Phospho-Stat3 (Tyr705) (#9145, CST, 1:2000), Stat3 (#9139, CST, 1:2000), IL6 (#12153, CST, 1:1000), β-Actin (8H10D10) (#3700, CST, 1:2000), GAPDH (#2118, CST, 1:10,000). Secondary antibodies were Anti-Mouse IgG Peroxidase Conjugate (A4416, Sigma, 1:5000), Anti-Rabbit IgG(H+L), Peroxidase conjugated (#31460, Thermo Scientific, 1:5000), Anti-Mouse IgG Peroxidase conjugated (1:5000), AntiGoat (sc-2354, Santa Cruz, 1:5000). Band intensities were quantified using ImageLab Version 6.1.0 (BioRad).

### Western Blot for secretory proteins

Equal numbers of cells were seeded in 35-mm dishes. Once the cell reaches 80% confluency, serum-free media was replaced and incubated for 48 to 72 hours. The conditioned media was then filtered with a 0.45-micron filter, and Trichloroacetic acid (Sisco, SRL) was added and incubated overnight. The secreted proteins were pelleted down and washed with acetone. Simultaneously, remaining cells were lysed directly with Laemmli buffer and were directly resolved on SDS-PAGE and processed for western blotting as described earlier. Ponceau S image was used as a loading control for the secretory proteins. β-actin was used to show equal loading of cells. Densitometric analysis was used to quantify the bands using ImageLab Version 6.1.0 (BioRad).

### Immunofluorescence and Immunohistochemistry

Cells were cultured on 50 μg/mL collagen-coated round coverslips in 24 plates. At the end of the culture period, cells were fixed with 4% (w/v) paraformaldehyde (Sigma-Aldrich Chemicals) followed by permeabilization with 0.1% Triton-X-100 (Sigma-Aldrich Chemicals). Primary antibodies of α-Smooth Muscle Actin (#19245, CST, 1:200), IGFBP5 (sc-515184, Santa Cruz, 1:200), IGFBP6 (1:200, AbCam), IGFBP7(1:200, SantaCruz). Secondary antibody conjugated with Alexa Fluor 568 (1:500, Invitrogen Bioservices) or Alexa Fluor 633 (1:500, Invitrogen Bioservices). F-actin was stained using TRITC-Phalloidin (1:200, Cat#P1951, Sigma-Aldrich Chemicals). Nuclei were stained using DAPI in PBS (1:1000, Sigma-Aldrich Chemicals). Slides were mounted using ProLong TM Glass Anti-Fade Mountant P36982, Invitrogen Bioservices) or Vecta Shield. The slides were visualized at 20X or 40X objective with Zeiss LSM 780 or Nikon AX Confocal Microscope. Images were processed using Fiji Software.

For tissue staining, thin sections from FFPE blocks were obtained on Poly-L-lysine-coated slides. The tissue sections were dewaxed and rehydrated in multiple changes of xylene and decreasing concentrations of ethanol. Antigen retrieval was performed by boiling in Tris EDTA (pH 9) and sodium citrate buffer (pH 6). These tissues were then blocked in horse serum and then incubated overnight with primary antibodies of α-Smooth Muscle Actin (#19245, CST, 1:500), IGFBP5 (sc-515184, Santa Cruz, 1:50), IGFBP6 (1:200, Cat# ab219560, Abcam) and IGFBP7 (1:200 dilution, Cat# ab171085, Abcam). Following thorough washing in PBST, secondary antibody conjugated with Alexa Fluor 488 goat anti-rabbit (Cat#A32731, Invitrogen, 1:500 dilution), Alexa Fluor 488 goat anti-mouse (Cat#A32723, Invitrogen, 1:500 dilution), Alexa Fluor 568 goat anti-mouse (Cat#A11031, Invitrogen, 1:500 dilution), Alexa Fluor 568 goat anti-rabbit (Cat#A11036, Invitrogen, 1:500 dilution), and Alexa Fluor 633 goat anti-mouse (Cat#A21052, Invitrogen, 1:500 dilution) was probed. Nuclei were stained using DAPI in PBS (1:1000 dilution, D9542, Sigma-Aldrich Chemicals). Slides were mounted using ProLong TM Glass Anti-Fade Mountant P36982, Invitrogen Bioservices) or Vecta Shield. The slides were visualised at 40X objective with Nikon AX Confocal Microscope. Images were processed using Fiji Software.

For immunohistochemistry, after primary antibody incubation, biotinylated antibodies were applied and incubated in Avidin-Biotin Complex Staining Kits (Cat#PK-6200, Vector Laboratories), followed by DAB staining and counterstaining of nuclei with hematoxylin. Dehydration in graded ethanol and xylene was performed before mounting. The slides were scanned with TissueScope iQ Slide Scanner at 0.25μm/pixel at 40X. The images were then processed and analysed using QuPath-0.5.1 (15) and Fiji.

### Multiplex immunofluorescence staining

A seven-colour multiplex immunofluorescence staining was performed using the OPAL^TM^ multiplexing method. The fixed tissues were cut into 4µm sections by Rotary Microtome (Leica RM2255, Germany). The sections were deparaffinized, rehydrated, subjected to heat-induced epitope retrieval and incubated with primary and secondary antibodies. The antibodies were visualised using an Opal 6-Plex Manual Detection Kit (NEL861001KT, Akoya Biosciences). The following primary antibodies at different dilutions were used: αSMA (ab5694, Abcam) at 1:100, FAP (EPR20021, ab207178, Abcam) at 1:75, and Pan-Keratin (#67306, Cell Signalling) at 1:50. IGFBP5 (SC-515184, Santa Cruz Biotechnology), IGFBP6 (ab219560, Abcam), and IGFBP7 (SC-365293, Abcam). Antibodies were visualised using the following tyramide dyes from the Opal Detection kit (NEL861001KT, Akoya Biosciences): Opal Polaris 480, Opal 520, Opal 570, Opal 620, Opal 690 and DIG-Opal 780. Nuclei were counterstained with Spectral DAPI (SKU FP1490, Akoya Biosciences). Sections were mounted with ProLong® Diamond Antifade Mountant (P36961, Thermo Fischer Scientific). Multiplex-stained slides were imaged using a PhenoImager Fusion system (Akoya Biosciences).

### siRNA-mediated knockdown

CAFs were transiently transfected with 20 nM of siRNAs (SMARTpool) targeting IGFBP5 (Cat# L-010897-00-0005), IGFBP6 (Cat#006625-00-0005) and IGFBP7 (Cat #L-008675-00-0005) or the control non-targeting siNT (Cat# D-001810-10-20, Dharamcon (ON-Target Plus SMART, Horizon). All siRNA transfections were performed using Dharmafect Transfection Reagent (Cat # T-2001-03) in OPTI-MEM (Invitrogen Bioservices) medium. After 6 hours, the medium was replaced with fresh RPMI containing 10% serum, and the cells were harvested for downstream experiments. The knockdown efficiency was validated by qRT-PCR or western blotting.

### Generation of stable knockdown cells

HEK293T cells were used to produce lentivirus and were maintained in DMEM media supplemented with HEPES buffer (HiMedia) and 10% heat-inactivated FBS. Lentiviral plasmids psPAX2 (Addgene, 12260), pMD2.G (Addgene, 12259), shIGFBP5 (TRCN0000062198), shIGFBP6 (TRCN0000062403), shIGFBP7 (TRCN0000077943), shControl (non-targeted control) (Cat# SHC002, Sigma-Aldrich) were transfected with Polyethyleneimine (Cat# 764892, Sigma-Aldrich). CAFs were seeded at 50% confluency and were infected with an appropriate amount of virus supernatant with 4 μg/mL polybrene (Cat# H9268-50G, Sigma-Aldrich Chemicals). After 48 hours, the cells were passaged and subjected to selection with hygromycin (Cat# ant-hg-1, Invivogen, France). Target gene knockdown efficiency was estimated using qRT-PCR and western blotting.

### Collagen contraction assay

The collagen contraction assay was performed as previously described (16). Briefly, CAFs (control/ knockdown cells) were embedded in rat tail Collagen-I (Cat#C9407, Corning) and incubated for 20 hours at 37⁰C. The gel was then imaged using BioRad Gel Doc, and the area of the contracted gel (in cm) was calculated using Fiji Software.

### Trans-well migration and invasion assay

As previously described (16), 8 μm trans-well (Cat#TCP257, HiMedia) was used for the migration and invasion assay. For invasion, the upper wells were coated with Matrigel (Growth Factor Reduced Basement Membrane Matrix, Cat#CMAPH000, Corning). Cells were seeded in the upper chamber in serum-free media. In the bottom chamber, complete media with EGF (Cat#PHG0311, Peprotech) (10 ng/mL) or conditioned media was added as a chemoattractant. After 16 hours, the wells were stained with crystal violet and non-migrated and non-invaded cells were wiped off thoroughly with a cotton swab. Four different fields from each well were captured using a Zeiss upright microscope. As there may be an overlap in some fields, cells were counted by averaging the results from four fields.

### Proliferation assay

mCherry-expressing H460 cells were seeded in a 96-well plate. Once the cells are adhered, the media was replaced with conditioned media derived from CAF (shControl or shIGFBPs) or treated with cisplatin (P4394-25MG, Sigma-Aldrich Chemicals). IC_50_ value of cisplatin for H460 cells was determined using MTT assay (Merck Life Sciences) and was found to be 4.8 μM. Also, the cisplatin-treated cells were subjected to live cell imaging (Sartorius Incucyte), where whole-well images were captured at 4X objective at an interval of 12 hours for a period of 3 days. The cell number was quantified using Basic Analyzer, Incucyte software, and the confluency percentage of each condition was plotted accordingly.

### Recombinant Protein Treatment

For recombinant protein treatment, fibroblast cells were seeded at 50-60% confluency. The next day, recombinant proteins, namely, TGFβ1 (10 ng/mL, Cat#100-21, Peprotech) and IL6 (10 ng/mL, Cat#200-06, Peprotech), were added to the media supplemented with 5% serum. When the indicated time points were reached, the cells were harvested for mRNA isolation or protein extraction.

### RNA Sequencing and Data Analysis

For RNA sequencing analysis, three independent batches of cells served as biological triplicates. Total RNA was isolated using Macherey-Nagel Nucleo Spin RNA Plus Kit. RNA library preparation and sequencing were performed (Medgenome, India). Alignment was performed using the STAR (v2.7.3a) aligner. Reads mapping to the ribosomal and mitochondrial genomes were removed before performing alignment. The raw read counts were estimated using featureCounts (v2.0.0). Read counts were normalised using DESeq2 to get the normalised counts. Log2FoldChange threshold was adjusted to 0.5. Heat map (Complex Heatmap) and volcano plot (EnhancedVolcano) of the differentially expressed genes were plotted using R programming (version 4.5.1). Pathway enrichment analysis was performed using ClusterProfiler (v4.0) (17). The upregulated and downregulated pathways were plotted based on the NES score; a positive NES score indicates upregulated pathways, while a negative NES score indicates downregulated pathways. Parallel to this, Gene Set Enrichment Analysis was performed using GSEA (GSEA v4.3.2, Human MSigDB v2023.1.Hs, Hallmark gene sets). CAF subtype gene signatures were downloaded from the Curated Cancer Cell Atlas (https://www.weizmann.ac.il/sites/3CA/) (18) and listed in the **Supplementary Table 2**. For the CAF subtype heatmap, only two biological replicates were considered, as one of the replicates was excluded due to the differences in clustering; still, validation of the genes was performed in more than three independent replicates. Enrichplot was used to visualise the enriched genes associated with the pathways. All enriched gene details and their pathways are uploaded in the **Supplementary Table 3 and 4.**

### NicheNet Analysis

NicheNet is a computational method that predicts ligand-target links between interacting cells (19). A set of potential ligands which are prevalent in the tumour microenvironment was defined based on the existing literature. To confirm that these ligands are also expressed in the lung tumour cells, we used our in-house RNA sequencing data of the tumour cells. Ligand-target matrix, ligand-receptor network and weighted networks were downloaded from NicheNet using R. To infer the ligand activity, differentially expressed genes between CAF siControl and siIGFBP5/6/7 were considered as the gene of interest, while all the other genes were considered as background genes. A ligand-target matrix was used with the defined potential ligands. The best upstream ligands whose receptors are expressed on the receiver cells were considered based on the AUPR score.

### In vivo tumorigenesis

All mice were procured from the in-house breeding facility (Laboratory Animal Facility, ACTREC). Luciferase and mCherry expressing H460 cells and GFP-expressing fibroblasts (shRNA mediated IGFBP5,6, 7 or shControl) were subcutaneously co-injected in the flank region (1:3 cell ratio). For these experiments, 6-week-old male NOD-SCID (NOD.Cg-Prkdcscid/J) mice were used. The tumour growth was monitored in vivo by injecting DLuciferin (IVISBRITE 122719, Perkin Elmer) and IVIS imaging (IVIS Spectrum or IVIS Lumina II). The images were processed using Living Image 4.8.2. All mice were examined every two days for health conditions and were sacrificed on the 15^th^ day by CO_2_ asphyxiation as per IAEC guidelines. Mice were sacrificed early to exclude the impact of the resident mouse fibroblasts. Tumour weight and volumes were measured after harvesting the tumours from the mice. Tumour volume was calculated as 0.5*(Length)*(Width)^2^.

### Survival Analysis

The gene profile and clinical data of lung adenocarcinoma were downloaded from TCGA using the GDC query. Counts were converted to log2CPM. For IGFBP5, gene sets with a log2 fold change less than 1 and an adjusted p-value below 0.05 were considered; while for IGFBP6 and IGFBP7, gene sets with a log2foldChange less than 1.5 with an adjusted p-value less than 0.05 were considered. The list of gene sets for calculating survival analysis is uploaded in the **Supplementary Table 5**. Overall survival was defined as the number of days from diagnosis to death. For patients with no information on death, the last date of follow-up was considered. Median score was calculated to determine high and low expression. KaplanMeier survival curves were plotted using the survival and survminer packages in R. The logrank test was used to determine statistical significance. The Cox regression model was used to determine the hazard ratio.

### Statistics and Reproducibility

Statistical analysis was performed using GraphPad Prism 8.0.2 software. Data are shown as mean ± SEM, unless otherwise indicated. Exact p-values are shown in the figures unless otherwise stated in the figure legends; p-values less than 0.05 are considered significant, and *ns* stands for non-significant. The Shapiro-Wilk test was used to assess the normality of the distributions (α=0.05). For data passing the normality test, a two-tailed unpaired Student’s ttest was used to determine the statistical significance between two groups, while for more than two groups, one-way or two-way ANOVA tests were used, and Dunnett’s multiple test correction was applied to compare means. For data that did not pass the normality test, a Kruskal-Wallis test was used, along with Dunn’s multiple comparison test where applicable. For patient samples, no statistical analysis was performed to determine the sample size; the samples included in the study are based on availability.

For animal experiments, Fig.5 h-j represents data from three independent experiments with similar results, with *n=4*-6 mice per group. All Western Blot experiments represent data from three independent experiments, except Supplementary Fig. 1b, which was conducted twice. All remaining in vitro experiments have been performed at least three times. All brightfield images are representative of four different fields taken from one experiment out of three independent experiments.

## Acknowledgements

We thank all the members of the Tumour Microenvironment Lab for their active discussions and support throughout the project. We thank the Digital Pathology Lab, Digital Imaging Facility, and Laboratory Animal Facility at ACTREC. We acknowledge Dr Sunil Shetty (ACTREC) for providing critical inputs, Nivedhya Venas and Dr Sabarinathan Radhakrishnan (NCBS, Bangalore) for reviewing our transcriptome analysis. We gratefully acknowledge Prof. Steven Johnsen and the Robert Bosch Center for Tumor Diseases (RBCT) for sharing the Akoya PhenoImager Fusion system. This work was supported by DST SERB-SRG (SRG/2021/001870-G) and intramural funding from ACTREC. G.D.Y. was funded by the ACTREC fellowship. We acknowledge the DBT-Ramalingaswami grant (BT/RLF/Reentry/09/2018), the Robert Bosch Stiftungand, The Capacity Building and Development of Novel Cutting Edge Research Activities grant (DPR 1/3(4)/2021/TMC/R&D-II/15063) by the Department of Atomic Energy, Govt. of India. The funders had no role in the study design, data collection and analysis, decision to publish or preparation of the manuscript.

## Authors’ Contributions

Conceptualization (G.D.Y., S.A.), Methodology (G.D.Y., J.S., M.D., K.N., S.R., R.T.), Data Curation (G.D.Y., K.N.), Formal Analysis (G.D.Y., S.A., S.Y., M.D.), Ethical clearance and patient data acquisition (K.P., R.K., S.Y., K.N., G.D.Y.), Funding Acquisition (S.A.), Writing-original draft (Y.G.D.), Writing-review & editing (G.D.Y., S.A.)

## Data Availability Statement

All data are shown in the article or Supplementary Information and Tables. TCGA-LUAD data were downloaded from the GDC portal (https://portal.gdc.cancer.gov/). Single-cell expression of lung studies was downloaded from the Curated Cancer Cell Atlas (https://www.weizmann.ac.il/sites/3CA/). All other raw data are available upon request from the corresponding authors.

## References

1. Lambrechts D, Wauters E, Boeckx B, Aibar S, Nittner D, Burton O, et al. Phenotype molding of stromal cells in the lung tumor microenvironment. Nat Med. 2018;24(8):1277–89.

2. Ohlund D, Handly-Santana A, Biffi G, Elyada E, Almeida AS, Ponz-Sarvise M, et al. Distinct populations of inflammatory fibroblasts and myofibroblasts in pancreatic cancer. J Exp Med. 2017;214(3):579–96.

3. Lavie D, Ben-Shmuel A, Erez N, Scherz-Shouval R. Cancer-associated fibroblasts in the single-cell era. Nat Cancer. 2022;3(7):793–807.

4. Huang H, Wang Z, Zhang Y, Pradhan RN, Ganguly D, Chandra R, et al. Mesothelial cell-derived antigen-presenting cancer-associated fibroblasts induce expansion of regulatory T cells in pancreatic cancer. Cancer Cell. 2022;40(6):656–73 e7.

5. Ozdemir BC, Pentcheva-Hoang T, Carstens JL, Zheng X, Wu CC, Simpson TR, et al. Depletion of carcinoma-associated fibroblasts and fibrosis induces immunosuppression and accelerates pancreas cancer with reduced survival. Cancer Cell. 2014;25(6):719–34.

6. Rhim AD, Oberstein PE, Thomas DH, Mirek ET, Palermo CF, Sastra SA, et al. Stromal elements act to restrain, rather than support, pancreatic ductal adenocarcinoma. Cancer Cell. 2014;25(6):735–47.

7. Baxter RC. IGF binding proteins in cancer: mechanistic and clinical insights. Nat Rev Cancer. 2014;14(5):329–41.

8. Forbes BE, McCarthy P, Norton RS. Insulin-like growth factor binding proteins: a structural perspective. Front Endocrinol (Lausanne). 2012;3:38.

9. Wang J, Ding N, Li Y, Cheng H, Wang D, Yang Q, et al. Insulin-like growth factor binding protein 5 (IGFBP5) functions as a tumor suppressor in human melanoma cells. Oncotarget. 2015;6(24):20636–49.

10. Sureshbabu A, Okajima H, Yamanaka D, Tonner E, Shastri S, Maycock J, et al. IGFBP5 induces cell adhesion, increases cell survival and inhibits cell migration in MCF-7 human breast cancer cells. J Cell Sci. 2012;125(Pt 7):1693–705.

11. Ding M, Bruick RK, Yu Y. Secreted IGFBP5 mediates mTORC1-dependent feedback inhibition of IGF-1 signalling. Nat Cell Biol. 2016;18(3):319–27.

12. Remsing Rix LL, Sumi NJ, Hu Q, Desai B, Bryant AT, Li X, et al. IGF-binding proteins secreted by cancer-associated fibroblasts induce context-dependent drug sensitization of lung cancer cells. Sci Signal. 2022;15(747):eabj5879.

13. Rupp C, Scherzer M, Rudisch A, Unger C, Haslinger C, Schweifer N, et al. IGFBP7, a novel tumor stroma marker, with growth-promoting effects in colon cancer through a paracrine tumor-stroma interaction. Oncogene. 2015;34(7):815–25.

14. Li D, Xia L, Huang P, Wang Z, Guo Q, Huang C, et al. Cancer-associated fibroblastsecreted IGFBP7 promotes gastric cancer by enhancing tumor associated macrophage infiltration via FGF2/FGFR1/PI3K/AKT axis. Cell Death Discov. 2023;9(1):17.

15. Bankhead P, Loughrey MB, Fernandez JA, Dombrowski Y, McArt DG, Dunne PD, et al. QuPath: Open source software for digital pathology image analysis. Sci Rep. 2017;7(1):16878.

16. Arandkar S, Furth N, Elisha Y, Nataraj NB, van der Kuip H, Yarden Y, et al. Altered p53 functionality in cancer-associated fibroblasts contributes to their cancer-supporting features. Proc Natl Acad Sci U S A. 2018;115(25):6410–5.

17. Wu T, Hu E, Xu S, Chen M, Guo P, Dai Z, et al. clusterProfiler 4.0: A universal enrichment tool for interpreting omics data. Innovation (Camb). 2021;2(3):100141.

18. Gavish A, Tyler M, Greenwald AC, Hoefflin R, Simkin D, Tschernichovsky R, et al. Hallmarks of transcriptional intratumour heterogeneity across a thousand tumours. Nature. 2023;618(7965):598–606.

19. Browaeys R, Saelens W, Saeys Y. NicheNet: modeling intercellular communication by linking ligands to target genes. Nat Methods. 2020;17(2):159–62.

20. Navab R, Strumpf D, Bandarchi B, Zhu CQ, Pintilie M, Ramnarine VR, et al. Prognostic gene-expression signature of carcinoma-associated fibroblasts in non-small cell lung cancer. Proc Natl Acad Sci U S A. 2011;108(17):7160–5.

21. Mathieson L, Koppensteiner L, Dorward DA, O’Connor RA, Akram AR. Cancerassociated fibroblasts expressing fibroblast activation protein and podoplanin in non-small cell lung cancer predict poor clinical outcome. Br J Cancer. 2024;130(11):1758–69.

22. Tyler M, Gavish A, Barbolin C, Tschernichovsky R, Hoefflin R, Mints M, et al. The Curated Cancer Cell Atlas provides a comprehensive characterization of tumors at singlecell resolution. Nat Cancer. 2025;6(6):1088–101.

23. Han Y, Wang Y, Dong X, Sun D, Liu Z, Yue J, et al. TISCH2: expanded datasets and new tools for single-cell transcriptome analyses of the tumor microenvironment. Nucleic Acids Res. 2023;51(D1):D1425–D31.

24. Ning Y, Schuller AG, Bradshaw S, Rotwein P, Ludwig T, Frystyk J, et al. Diminished growth and enhanced glucose metabolism in triple knockout mice containing mutations of insulin-like growth factor binding protein-3, -4, and -5. Mol Endocrinol. 2006;20(9):2173–86.

25. Kato T, Jenkins RP, Derzsi S, Tozluoglu M, Rullan A, Hooper S, et al. Interplay of adherens junctions and matrix proteolysis determines the invasive pattern and growth of squamous cell carcinoma. Elife. 2023;12.

26. Cumming J, Maneshi P, Dongre M, Alsaed T, Dehghan-Nayeri MJ, Ling A, et al. Dissecting FAP+ Cell Diversity in Pancreatic Cancer Uncovers an Interferon-Response Subtype of Cancer-Associated Fibroblasts with Tumor-Restraining Properties. Cancer Res. 2025;85(13):2388–411.

27. Hu H, Piotrowska Z, Hare PJ, Chen H, Mulvey HE, Mayfield A, et al. Three subtypes of lung cancer fibroblasts define distinct therapeutic paradigms. Cancer Cell. 2021;39(11):1531–47 e10.

28. Dorris ER, Tazzyman SJ, Moylett J, Ramamoorthi N, Hackney J, Townsend M, et al. The Autoimmune Susceptibility Gene C5orf30 Regulates Macrophage-Mediated Resolution of Inflammation. J Immunol. 2019;202(4):1069–78.

29. Broz MT, Ko EY, Ishaya K, Xiao J, De Simone M, Hoi XP, et al. Metabolic targeting of cancer associated fibroblasts overcomes T-cell exclusion and chemoresistance in soft-tissue sarcomas. Nat Commun. 2024;15(1):2498.

30. Mantel I, Sadiq BA, Blander JM. Spotlight on TAP and its vital role in antigen presentation and cross-presentation. Mol Immunol. 2022;142:105–19.

31. Li L, Ugalde AP, Scheele C, Dieter SM, Nagel R, Ma J, et al. A comprehensive enhancer screen identifies TRAM2 as a key and novel mediator of YAP oncogenesis. Genome Biol. 2021;22(1):54.

32. Chhabra Y, Weeraratna AT. Fibroblasts in cancer: Unity in heterogeneity. Cell. 2023;186(8):1580–609.

33. Ma C, Yang C, Peng A, Sun T, Ji X, Mi J, et al. Pan-cancer spatially resolved singlecell analysis reveals the crosstalk between cancer-associated fibroblasts and tumor microenvironment. Mol Cancer. 2023;22(1):170.

34. Rousse S, Lallemand F, Montarras D, Pinset C, Mazars A, Prunier C, et al. Transforming growth factor-beta inhibition of insulin-like growth factor-binding protein-5 synthesis in skeletal muscle cells involves a c-Jun N-terminal kinase-dependent pathway. J Biol Chem. 2001;276(50):46961–7.

35. Kojima H, Kunimoto H, Inoue T, Nakajima K. The STAT3-IGFBP5 axis is critical for IL-6/gp130-induced premature senescence in human fibroblasts. Cell Cycle. 2012;11(4):730–9.

36. Biffi G, Oni TE, Spielman B, Hao Y, Elyada E, Park Y, et al. IL1-Induced JAK/STAT Signaling Is Antagonized by TGFbeta to Shape CAF Heterogeneity in Pancreatic Ductal Adenocarcinoma. Cancer Discov. 2019;9(2):282–301.

37. Badia IMP, Velez Santiago J, Braunger J, Geiss C, Dimitrov D, Muller-Dott S, et al. decoupleR: ensemble of computational methods to infer biological activities from omics data. Bioinform Adv. 2022;2(1):vbac016.

38. Long X, Xiong W, Zeng X, Qi L, Cai Y, Mo M, et al. Cancer-associated fibroblasts promote cisplatin resistance in bladder cancer cells by increasing IGF-1/ERbeta/Bcl-2 signalling. Cell Death Dis. 2019;10(5):375.

39. Tao L, Huang G, Wang R, Pan Y, He Z, Chu X, et al. Cancer-associated fibroblasts treated with cisplatin facilitates chemoresistance of lung adenocarcinoma through IL-11/IL-11R/STAT3 signaling pathway. Sci Rep. 2016;6:38408.

40. Linares J, Sallent-Aragay A, Badia-Ramentol J, Recort-Bascuas A, Mendez A, Manero-Ruperez N, et al. Long-term platinum-based drug accumulation in cancerassociated fibroblasts promotes colorectal cancer progression and resistance to therapy. Nat Commun. 2023;14(1):746.

41. Hanley CJ, Mellone M, Ford K, Thirdborough SM, Mellows T, Frampton SJ, et al. Targeting the Myofibroblastic Cancer-Associated Fibroblast Phenotype Through Inhibition of NOX4. J Natl Cancer Inst. 2018;110(1):109–20.

42. Herzog BH, Baer JM, Borcherding N, Kingston NL, Belle JI, Knolhoff BL, et al. Tumor-associated fibrosis impairs immune surveillance and response to immune checkpoint blockade in non-small cell lung cancer. Sci Transl Med. 2023;15(699):eadh8005.

43. Chen Y, McAndrews KM, Kalluri R. Clinical and therapeutic relevance of cancerassociated fibroblasts. Nat Rev Clin Oncol. 2021;18(12):792–804.

44. Galbo PM, Jr., Zang X, Zheng D. Molecular Features of Cancer-associated Fibroblast Subtypes and their Implication on Cancer Pathogenesis, Prognosis, and Immunotherapy Resistance. Clin Cancer Res. 2021;27(9):2636–47.

45. Li K, Wang R, Liu GW, Peng ZY, Wang JC, Xiao GD, et al. Refining the optimal CAF cluster marker for predicting TME-dependent survival expectancy and treatment benefits in NSCLC patients. Sci Rep. 2024;14(1):16766.

46. Baxter RC. Signaling Pathways of the Insulin-like Growth Factor Binding Proteins. Endocr Rev. 2023;44(5):753–78.

47. Su M, Zhao W, Jiang H, Zhao Y, Liao Z, Liu Z, et al. Endothelial IGFBP6 suppresses vascular inflammation and atherosclerosis. Nat Cardiovasc Res. 2025;4(2):145–62.

48. Nikulin S, Razumovskaya A, Poloznikov A, Zakharova G, Alekseev B, Tonevitsky A. ELOVL5 and IGFBP6 genes modulate sensitivity of breast cancer cells to ferroptosis. Front Mol Biosci. 2023;10:1075704.

49. Iosef C, Vilk G, Gkourasas T, Lee KJ, Chen BP, Fu P, et al. Insulin-like growth factor binding protein-6 (IGFBP-6) interacts with DNA-end binding protein Ku80 to regulate cell fate. Cell Signal. 2010;22(7):1033–43.

50. Broad RV, Jones SJ, Teske MC, Wastall LM, Hanby AM, Thorne JL, et al. Inhibition of interferon-signalling halts cancer-associated fibroblast-dependent protection of breast cancer cells from chemotherapy. Br J Cancer. 2021;124(6):1110–20.

51. Cao Z, Meng Z, Li J, Tian Y, Lu L, Wang A, et al. Interferon-gamma-stimulated antigen-presenting cancer-associated fibroblasts hinder neoadjuvant chemoimmunotherapy efficacy in lung cancer. Cell Rep Med. 2025;6(3):102017.

52. Huang L, Lu W, Wu R, Li Y, Ou Z, Chen J, et al. Interferon-driven CAF reprogramming augments immunogenic response to neoadjuvant radiotherapy in colorectal cancer. Cell Rep Med. 2025;6(8):102251.

53. Sahai E, Astsaturov I, Cukierman E, DeNardo DG, Egeblad M, Evans RM, et al. A framework for advancing our understanding of cancer-associated fibroblasts. Nat Rev Cancer. 2020;20(3):174–86.

54. Chrisochoidou Y, Roy R, Farahmand P, Gonzalez G, Doig J, Krasny L, et al. Crosstalk with lung fibroblasts shapes the growth and therapeutic response of mesothelioma cells. Cell Death Dis. 2023;14(11):725.

