## Supplementary Figures for "IGFBPs define distinct pro-tumorigenic CAF subtypes in lung cancer tumour-microenvironment"

1

2 # Supplementary Figures

Supplementary Fig. 1

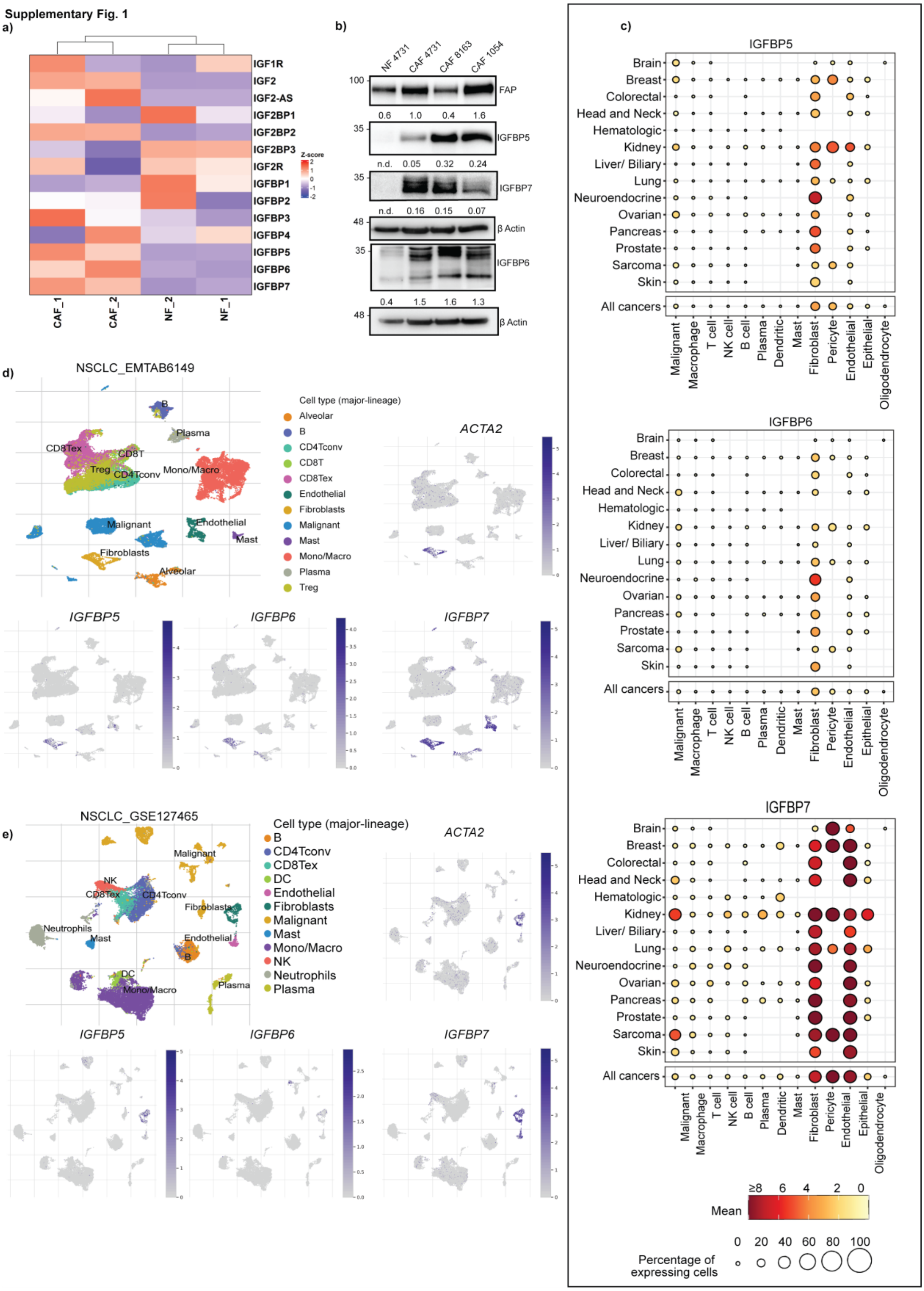


3

**Supplementary Fig. 1 Lung cancer-associated fibroblasts exhibit higher IGFBP5, 6 and 7 expressions. a)** Heatmap showing differentially expressed genes between lung cancer patient-derived NF and CAF, data taken from Arandkar *et al.*, 2018. Vertical rows indicate different samples with two biological replicates each, horizontal rows indicate genes belonging to the IGFBP family, coloured according to log-transformed transcripts in z-score units (Red: High expression, blue: low expression), **b)** Immunoblots showing IGFBP5, 6 and 7 expressions in different lung cancer patient-derived fibroblast cell lines (4731, 8163, 1054). β-Actin served as the loading control. Densitometric analysis was done using Image Lab (v6.1.0), n= 3 biological replicates, **c)** Dot plot representing IGFBP5 (top), IGFBP6 (middle) and IGFBP7 (bottom) expressions in different cell types across all cancers. Data was generated from the Curated Cancer Cell Atlas. The size of the circle dot represents the percentage of expressing cells, while the colour represents the mean expression level, as indicated in the scale bar (below), **d)** UMAP plots showing ACTA2, IGFBP5,6 and 7 expressions in fibroblasts in two publicly available NSCLC single-cell RNA sequencing datasets (NSCLC_EMTAB6149 and NSCLC_GSE127465). Data was generated from Tumour Immune Single-Cell Hub2 (TISCH2).

Supplementary Fig.2

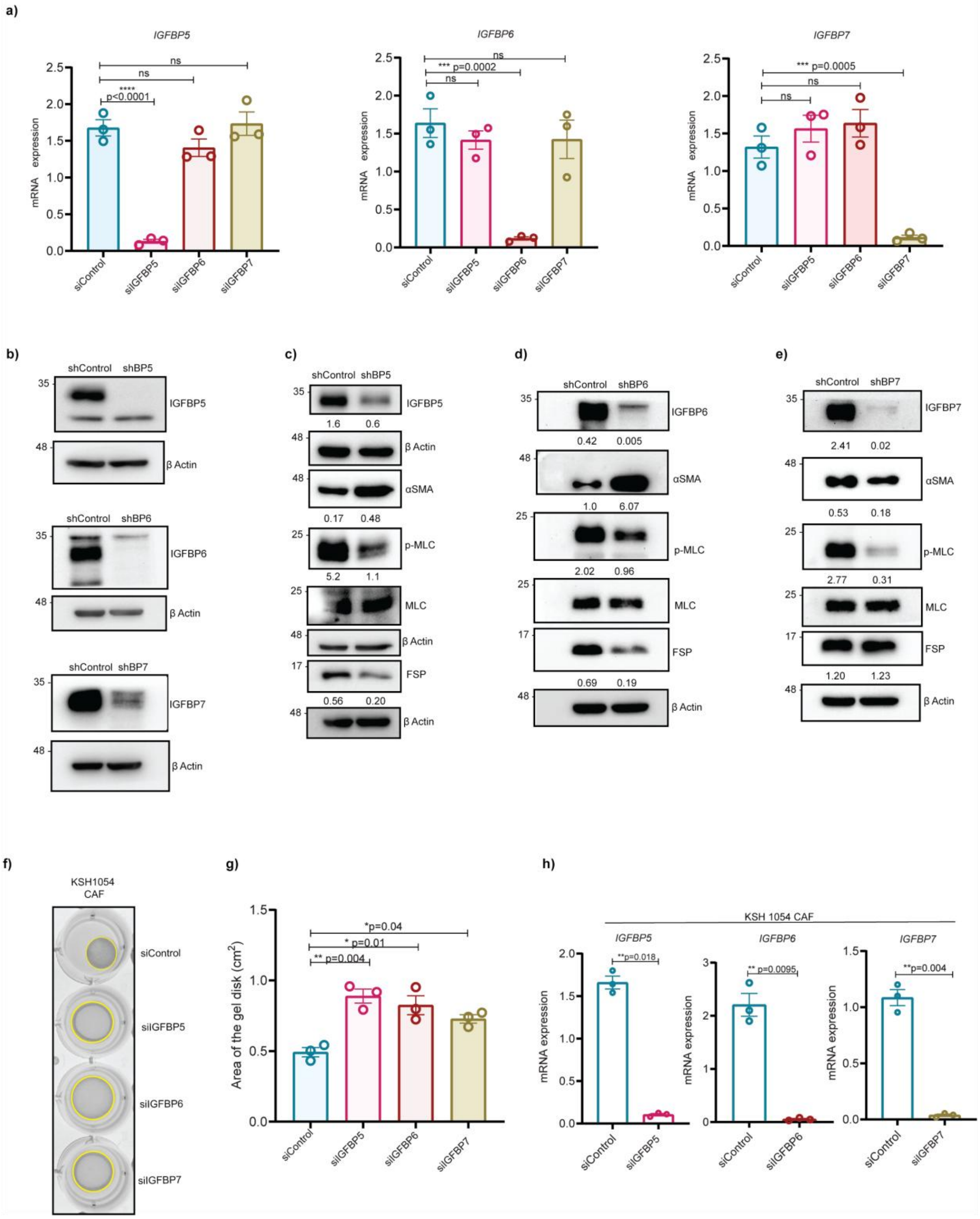


**Supplementary Fig. 2 IGFBP5, 6 and 7 regulate CAF's properties. a)** mRNA expression of IGFBP5, 6 and 7 upon individual siRNA-mediated knockdown in CAF 4731. Data represent mean ± SEM, n=3. **b)** Validation of IGFBP5, 6 and 7 knockdowns in stable knockdown CAF cells. β-Actin served as the loading control, **c), d) & e)** Immunoblots representing p-MLC, FSP and αSMA levels upon individual IGFBP knockdown. β-Actin served as the loading control. Densitometric analysis was done using Image Lab **f)** Image representing collagen contraction upon individual knockdown of IGFBP5, 6 and 7 in KSH1054 CAF cells. Yellow circles mark the boundary of the gel, **g)** Quantification of the area of the gel disk is represented as a bar plot with mean ± SEM, n=3. **h**) Knockdown validation of IGFBP5, 6 and 7 at the mRNA level in KSH1054 cells. Data represent mean ± SEM, n=3. To determine statistical significance, an unpaired t-test with Welch's correction was used; exact p-values are indicated in the graph. **(a & g)** To determine statistical significance, One-way ANOVA with Dunnett's multiple comparison was used; exact p-values are indicated in the graph.

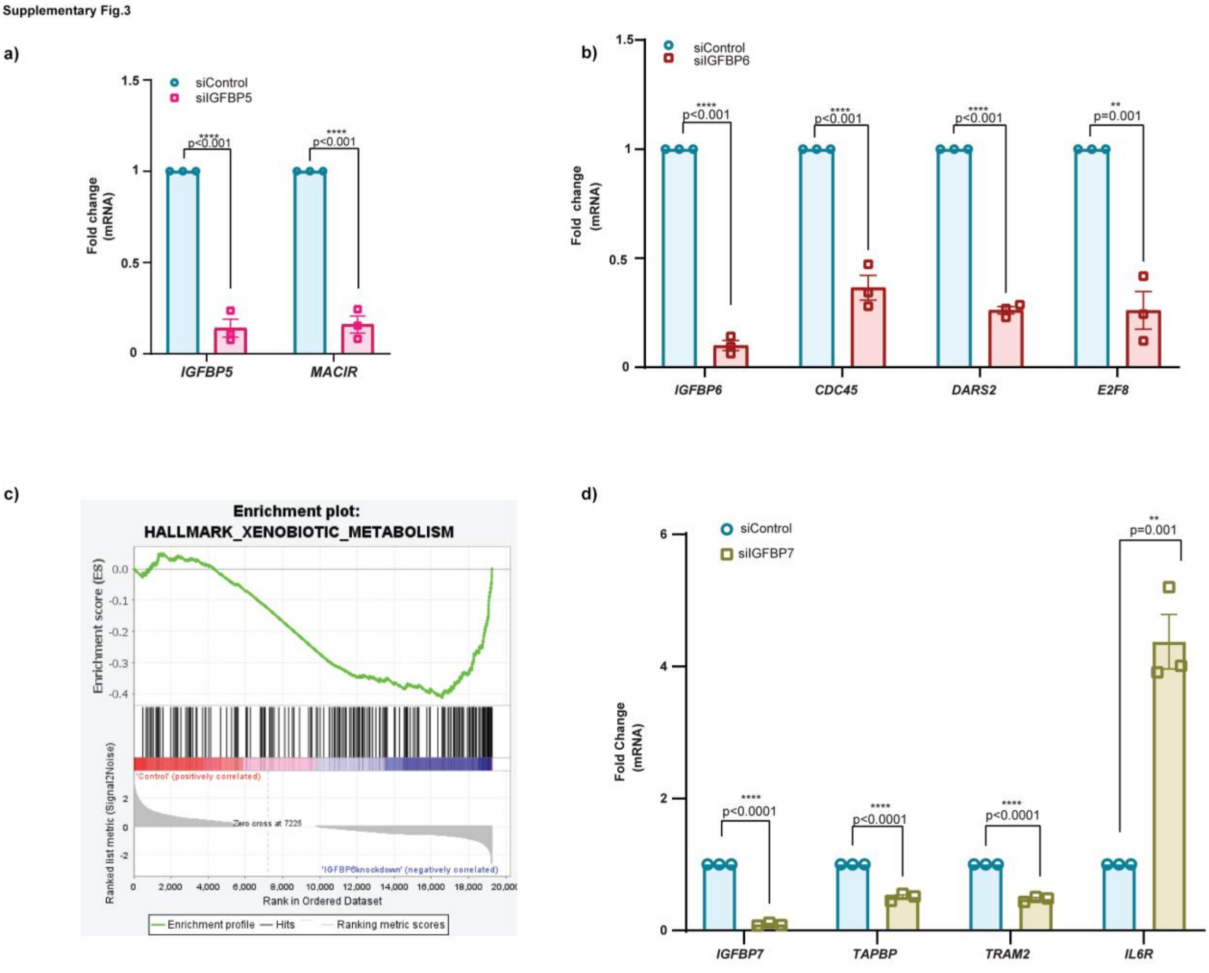


**Supplementary Fig.3: Validation of genes after CAF siIGFBP knockdown. a)** Fold-change mRNA expression of the *MACIR* gene using qRT-PCR in siControl versus siIGFBP5. **b)** Fold-change mRNA expression of cell-cycle related genes (*CDC45, DARS2, E2F8*) in siControl versus siIGFBP6. **c)** GSEA plot of “HALLMARK_XENOBIOTIC_METABOLISM” in siControl versus siIGFBP6. **d)** Fold-change mRNA expression of TAPBP, TRAM2, IL6R using qRT-PCR in siControl versus siIGFBP7. **(a, b, d)** Data represent mean ± SEM of three biological replicates. To determine statistical significance, an unpaired t-test was used; the p-value is indicated in the graph.

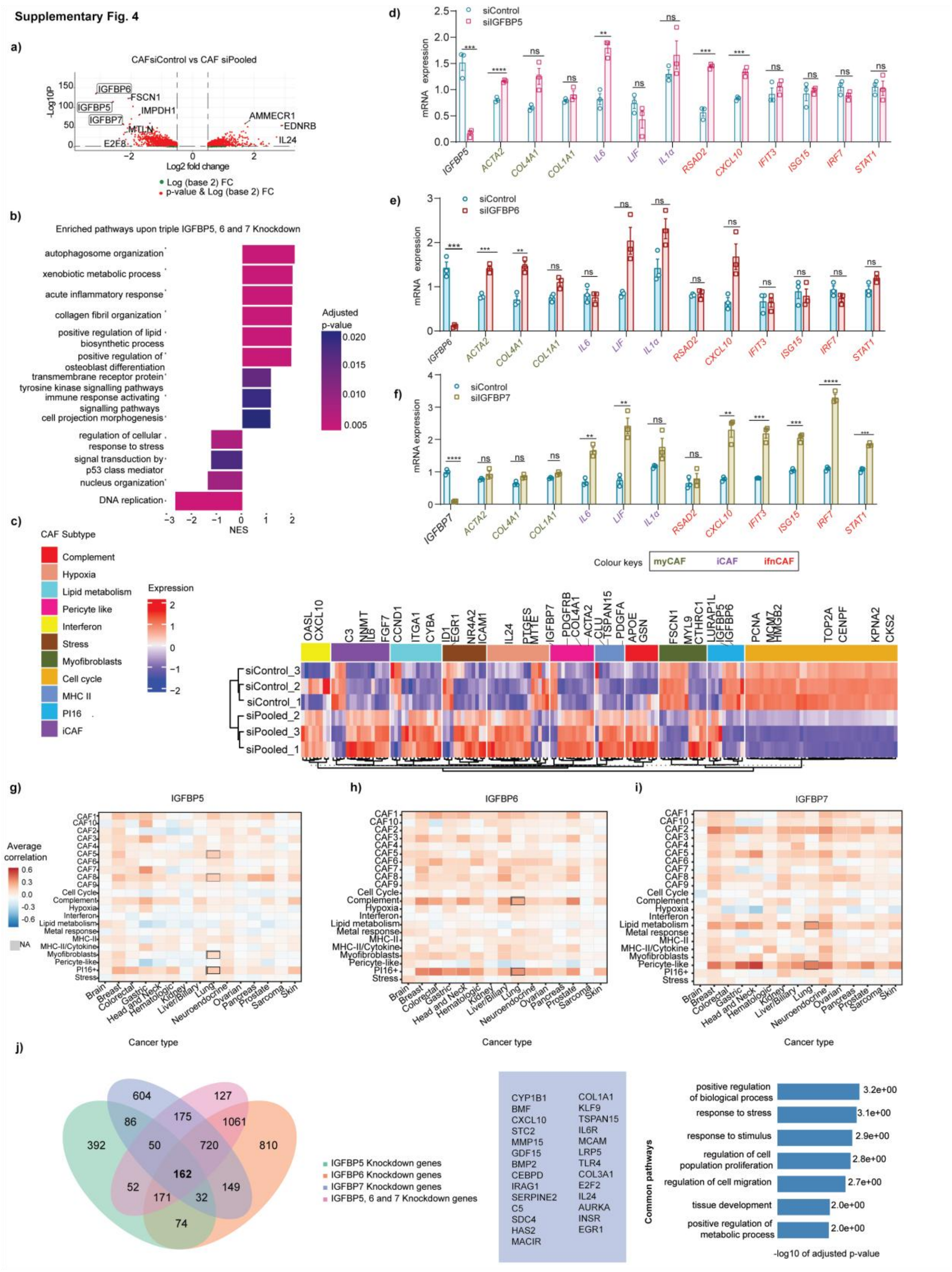


**Supplementary Fig. 4 CAF subtype gene validation in individual IGFBP knockdowns and RNA-sequencing analysis of Pooled (IGFBP5, 6 and 7) knockdown**. **a)** Volcano plot representing differentially expressed genes between CAF siNT (non-targeted control) vs siPooled (n=3 biological replicates) **b)** Enriched pathways based on the differentially expressed genes upon Pooled knockdown **c)** Heatmap comparing the published CAF subtype gene sets with CAF siNT versus siPooled **d, e & f)** RT-qPCR of myofibroblastic (myCAF), inflammatory (iCAF) and interferon-response related (ifn) CAF genes upon IGFBP5, 6 and 7 knockdown, respectively. Data represent mean ± SEM of three biological replicates. **g, h & i)** Heatmaps showing, for fibroblasts, the average Pearson correlation of IGFBP5, 6 and 7, respectively, with meta-program scores in each cancer type. NAs indicate insufficient data or zero variance. **j)** Venn diagram showing common genes between individual and Pooled knockdown of IGFBP5, 6 and 7 (left), list of common differentially expressed genes between individual and pooled knockdowns (middle) and pathways enriched with the common genes regulated by IGFBP5, 6 and 7 (right)

**Supplementary Fig. 5**

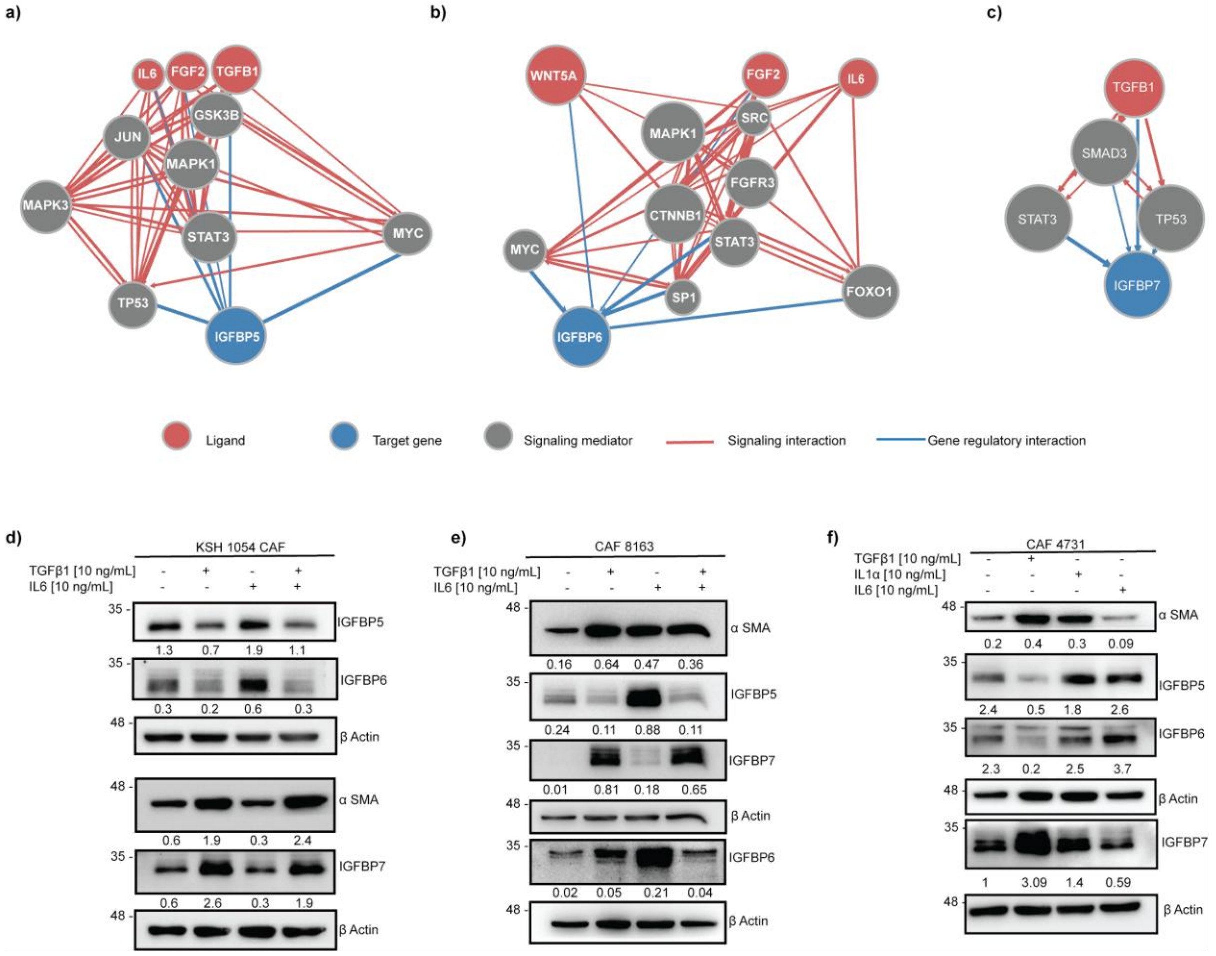

**Supplementary Fig. 5 TGFβ1 and IL6 regulate IGFBP expression in CAFs. a-c)** Network of interactions from NicheNet scores between ligand, signalling mediators, transcriptional regulators, and target genes (IGFBP5 **(a)**, IGFBP6 **(b)**and IGFBP7 **(c)**). The ligand node is indicated in red, the target gene nodes in blue and the nodes of signalling mediators in grey. Red-coloured edges represent signalling interactions, while blue-coloured edges represent gene regulatory interactions. **(d-f)** Immunoblots representing IGFBP5, 6 and 7 expressions in KSH 105 **(d),** CAF 8163 **(e),** and CAF 4731 **(f)** after treatment with TGFβ1 [10ng/mL], IL- 1α [10 ng/mL] and IL6 [10ng/mL] for 48 hours. Densitometric analysis was performed using Image Lab (v6.1.0), normalised to their corresponding β Actin.

Supplementary Fig. 6

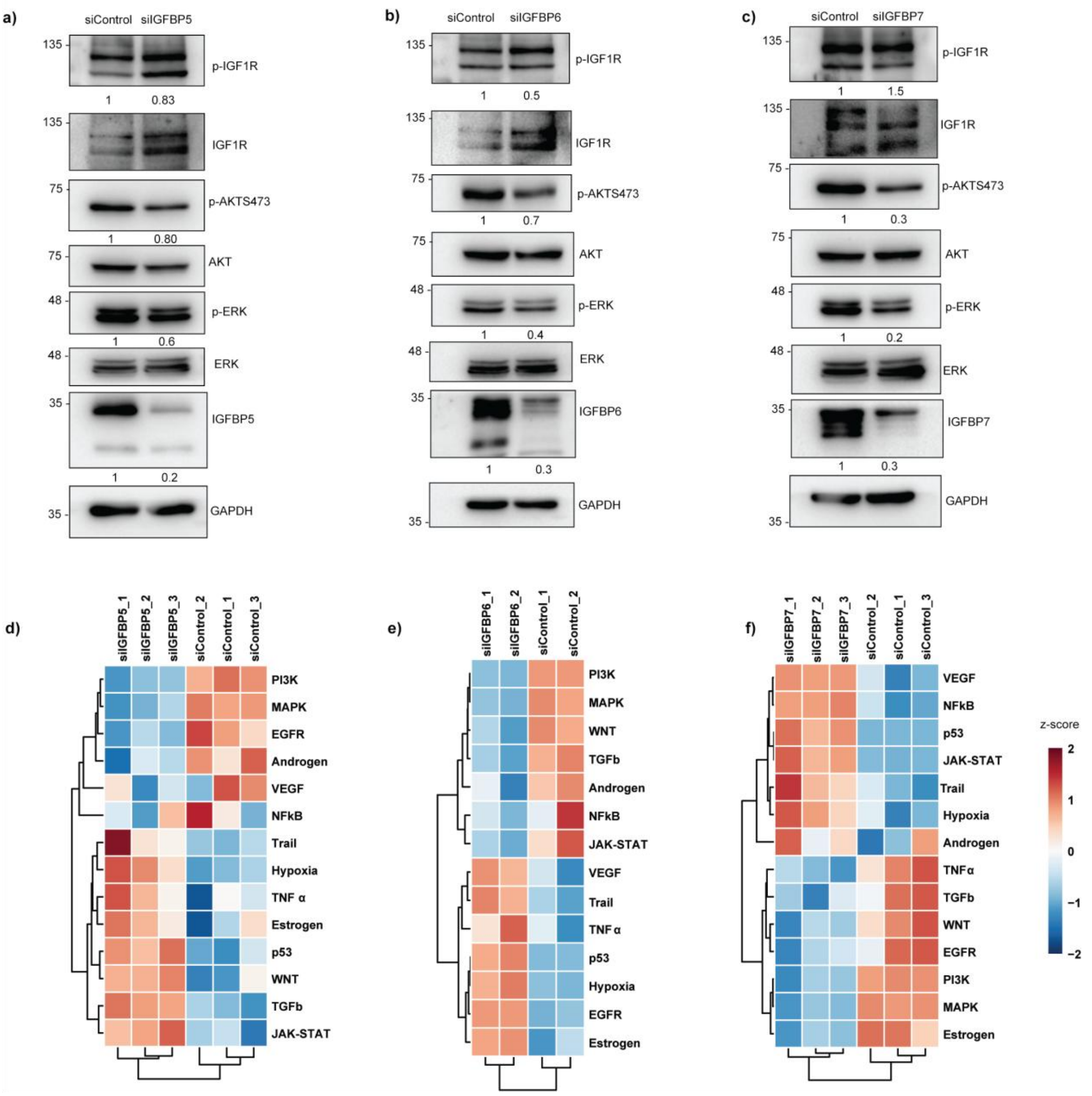


**Supplementary Fig. 6 IGFBP5, 6 and 7 exhibit IGF-independent pathways in CAFs. a, b & c)** Immunoblots representing IGF-signalling pathways upon individual IGFBP knockdowns in CAF. Densitometric analysis was done using Image Lab (v6.1.0). **d, e & f)** Pathway activity inference of CAF siControl and siIGFBP5, 6 and 7 showing different pathway scores using decoupleR.

Supplementary Fig. 7

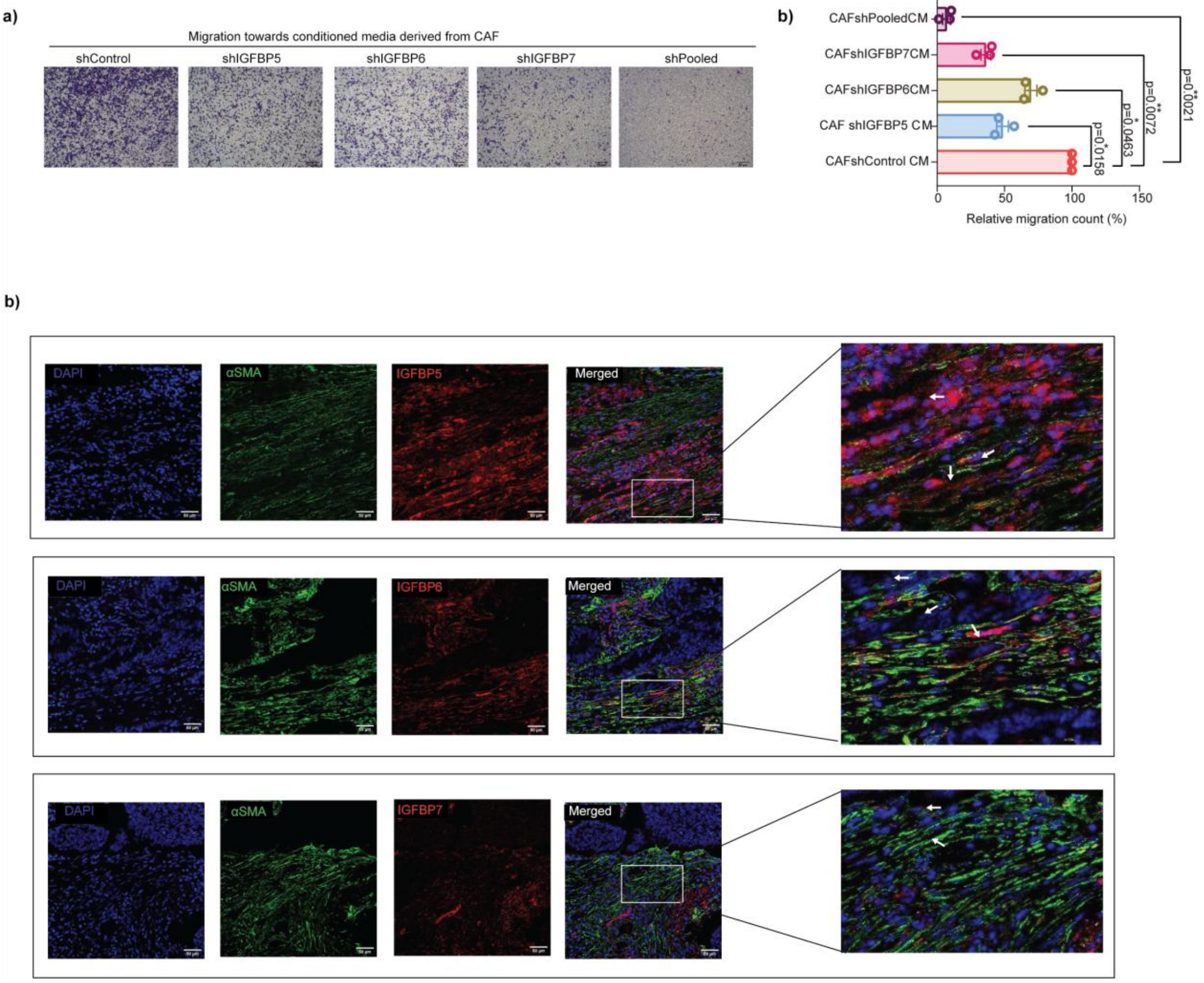


**Supplementary Fig. 7. IGFBP5, 6, and 7-expressing CAFs impact tumour cell migration.** **a)** Representative brightfield images of migrated cells (crystal violet stained) of tumour cells towards conditioned media (CM) derived from individual and pooled IGFBP5, 6 and 7 knockdowns in CAF. Images were captured 16 hours after the non-migrated cells in the upper chamber were wiped off. **b)** Cell count of the migrated cells is taken as an average of four different fields of one experiment, data represents mean (relative count with respect to CAF shControl (%) ± SEM, n=3 biological replicates. To determine statistical significance, One-way ANOVA with Dunnett's multiple comparison was used; exact p-values are indicated in the graph. **c)** Immunofluorescence dual staining of αSMA (green), IGFBP5/6/7 (red), DAPI (blue) from different fields of a patient case. The arrow indicates the co-expression of αSMA and respective IGFBP5/6/7: scale bar, 50 μm.
